# How to be dispensable: genomic and transcriptomic determinants in maize genes

**DOI:** 10.64898/2026.04.25.720561

**Authors:** Johann Joets, Maëva Mollion, Kevin Baudry, Maud Fagny, Olivier Turc, Llorenç Cabrera-Bosquet, Sylvie Coursol, Claude Welcker, Peter Rogowsky, Harry Belcram, Agnès Rousselet, Anthony Venon, Sandrine Chaignon, Stéphanie Pateyron, Jérôme Laplaige, Christine Paysant Le Roux, Véronique Brunaud, Marie-Laure Martin, Carine Palaffre, William Marande, Maud I. Tenaillon, Clémentine Vitte

## Abstract

**Background:** Plant genomes harbour a substantial proportion of dispensable genes – present only in a subset of individuals – that differ from ubiquitously shared core genes in multiple genomic and expression features. While these differences have been repeatedly documented, the factors shaping gene dispensability remain poorly understood.

**Results:** We assembled a pan-gene set from eight maize inbred lines from American and European germplasms, together with their transcriptomic profile across 22 tissues/conditions, revealing the genomic and transcriptomic determinants of maize gene dispensability. Multivariate analysis demonstrates that gene expression level and purifying selection – rather than gene size – are the primary factors distinguishing core from dispensable genes. Dispensable genes overlap *Helitrons* at 4.6 times the rate of core genes, implicating *Helitron*-mediated gene capture as a major mechanism of dispensable gene formation. Classifying genes into stably expressed, variably expressed, and on-off categories shows that all three classes contain dispensable genes, though in different proportions than for core genes. Contrary to previous assumptions, we show that dispensable genes can participate in basal biological functions just as core genes, and that gene duplication likely provides only a partial mechanism for functional complementation of accessory genes absence.

**Conclusions:** Our results provide novel insights into the molecular and evolutionary factors distinguishing core from dispensable genes and into the biological mechanisms shaping gene dispensability in maize, and demonstrate that classifying genes by transcriptional patterns provides a powerful framework for understanding the biological functions and evolutionary dynamics of both core and dispensable genes.

## Background

Plant genomes harbour a substantial proportion of genes present only in a subset of individuals within a species – 40% in *Arabidopsis thaliana*^1^, 45% in maize^2^, 54% in *Brassica rapa*^3^, 60% in tomato^4^, 70% in rice^5^ and 75% in *Brachypodium distachyon*^6^. Unlike core genes – which are conserved across all members of a species and essential for basic survival and reproduction – ‘accessory’ or ‘dispensable’ genes are variably present and not strictly required under all conditions. Their presence may nevertheless confer a fitness advantage to individuals under specific environmental conditions^7–9^.

In plants, core and dispensable genes differ markedly across a range of molecular and evolutionary features. Compared to core genes, dispensable genes display, on average, a higher 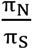 ratio (the ratio of non-synonymous to synonymous nucleotide substitutions)^10^, indicative of faster evolution and/or relaxed purifying selection. They are also shorter, expressed under fewer conditions, and exhibit lower expression levels. These characteristics make functional annotation of dispensable genes particularly challenging; consequently, they remain less well characterised than core genes. Nonetheless, Gene Ontology analyses consistently reveal enrichment in functions related to environmental responses and developmental processes, rather than in basal biological functions^6,11–13^ – a pattern consistent with a primary role in adaptive processes enabling organisms to respond to environmental challenges^14^.

Despite these well-documented differences, the interrelationships among these features – gene size, expression level,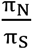 Effective Number of Codons (ENC), and biological function – and the mechanisms by which they collectively shape gene dispensability remain poorly understood. Beyond standard GO enrichment analyses, which offer only a broad overview of gene function, few studies have exploited the full range of molecular features of accessory genes to refine our understanding of their biology. In this regard, a negative correlation between gene size and the frequency of structural variants has been reported in several plant species^15–19^, suggesting that a mechanism favoring shorter structural variants may contribute to the size bias observed for dispensable genes. The dynamics of transposable elements – and of *Helitrons* in particular – likely also play a role in shaping the dispensable genome^20^. In maize, 90% of *Helitrons* carry at least one gene fragment^21^; while *Helitrons* can sequentially capture multiple fragments, most carry only one or two^21^, potentially generating transcripts shorter than the source gene copies and thus contributing to the gene size bias of the dispensable genome.

Here, we constructed a pan-gene set for eight maize inbred lines and quantified gene expression across 15 tissues and six developmental stages, including seven tissues sampled under two contrasting water regimes. Joint analysis of the genomic, evolutionary, and expression features of this pan-gene set provides new insights into the factors distinguishing core from dispensable genes and into the biological mechanisms shaping gene dispensability in maize. Our findings indicate that dispensable gene compartment is primarily driven by gene expression level and purifying selection rather than gene size, and highlight the role of *Helitrons* in dispensable gene dynamics. Contrary to previous reports, we also provide evidence for the involvement of dispensable genes in basal biological functions.

## Results

### Generation of a high-quality genome sequence assembly and annotation

To characterise the transcriptional and functional properties of core and dispensable genes, we generated genome sequence assemblies for seven maize lines representing American and European genetic groups and transcriptomes for these seven maize lines and B73. They were chosen to cover a wide range of the genetic diversity that contributed to maize elite material currently cultivated in Europe (Supplementary table 1). Genome assemblies were produced following the approach described in ^22^, and showed high completeness, with an average read self-mapping rate of 99.53% (Supplementary table 2). As expected, chromosomes from genetically closer pairs of genotypes exhibited the greatest sequence contiguity (Supplementary figure 1). BUSCO scores are similar to these of B73 NAM5 confirming the high quality of the gene space in our assemblies (Supplementary figure 2).

To support both genome annotation and transcriptional analysis of core and dispensable genes across these seven lines and B73, we generated an extensive mRNA-seq dataset comprising three biological replicates for each of 15 tissues sampled across six developmental stages, five of which were collected under two contrasting water regimes (Supplementary table 6). Gene models were deliberately not filtered on predicted protein length, thus ensuring retention of genes encoding short proteins. This approach yielded between 60,783 and 63,606 predicted gene models per genotype (Supplementary table 3).

### Transcriptional classes distributions reveal differences between core and dispensable genes

To characterise the extent of gene structural variation across the eight genotypes, we assembled a pan-gene set following the workflow outlined in Supplementary figure 3. Briefly, genes present in all eight genotypes were classified as ‘core’, while genes absent from at least one genotype were classified as ‘dispensable’. The pan-gene set across the eight lines comprised 76,358 unique gene models, of which 46,163 (60%) were core. Each genotype harboured between 14,620 and 17,443 dispensable genes (24.05% to 27.42% of total genes) (Supplementary table 3).

We compared core and dispensable genes across several genomic, molecular and evolutionary features: gene length and content (gene and protein size, GC% of the CDS sequence), evolutionary rate 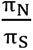 and expression-related variables (Effective Number of Codon, expression level and expression breadth). As expected^14^, the two gene sets differed significantly: dispensable genes displayed lower expression levels, smaller gene size and lower GC% but higher 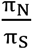 and ENC than core genes (Figure 1A-E). These differences are consistent with relaxed purifying selection and weaker codon usage bias in dispensable genes. Dispensable genes also showed more tissue- and condition-specific expression than core genes (Figure 1F). To gain further insights into the functional characteristics of core and dispensable genes, we classified genes into three categories based on expression patterns: Stably Expressed Genes (SEGs, expressed in all samples with CV ≤ 0.1), Variably Expressed Genes (VEGs, expressed in all samples with CV > 0.1, and On-Off Genes (OOGs, expressed in a subset of samples only).

**Figure 1:**
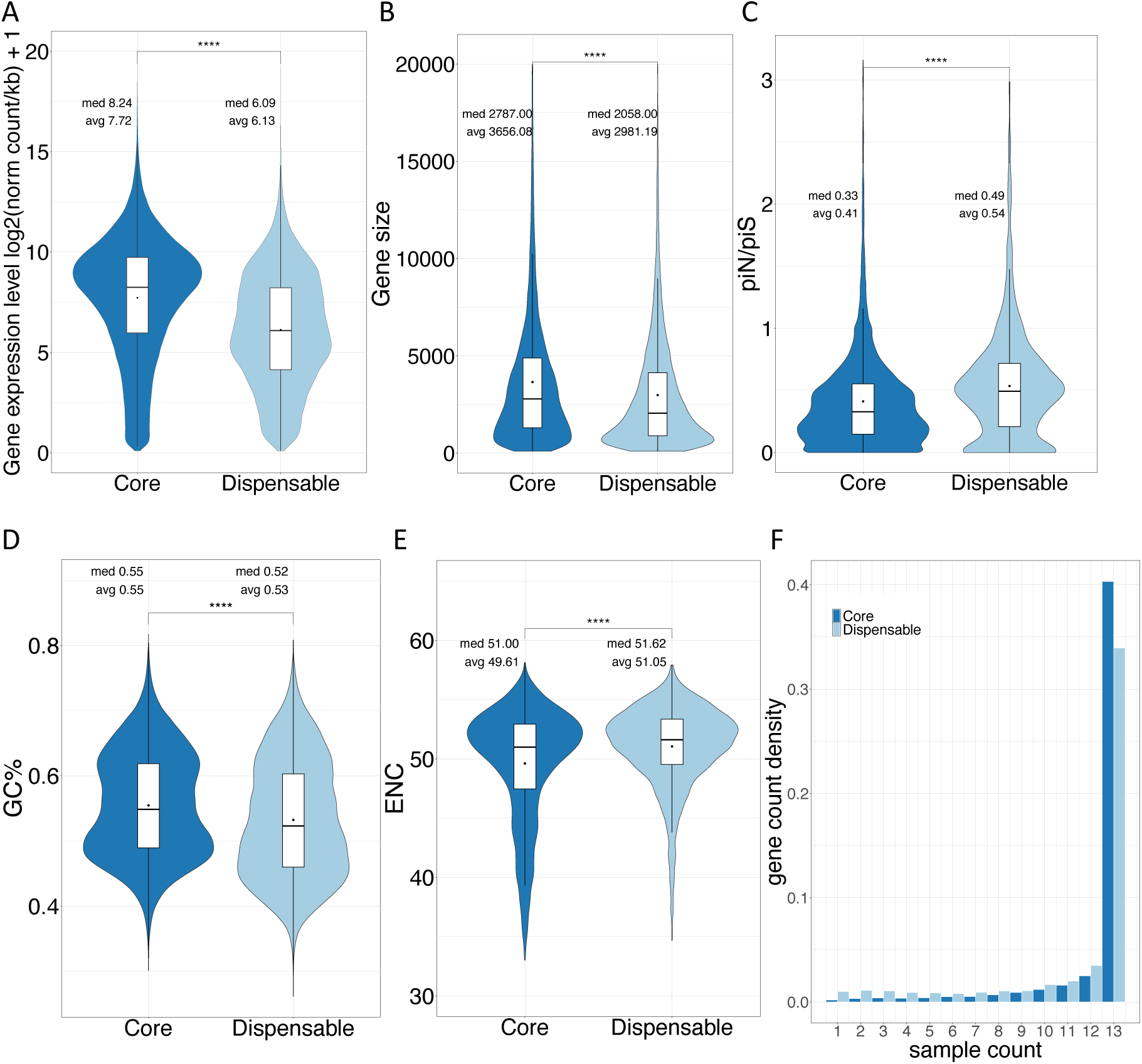
Features of core and dispensable genes. Violin plots for gene size (A), expression level (B),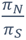 (*C*), Proportion of GC in coding sequence (D), ENC (E), Expression breadth excluding water deficit condition (F). Showed results were obtained on the B73 genotype and are similar for all the other genotypes under study (not shown). (A-D) p-values were obtained using a t-test. All p-values were lower than 0.001(****). Med = median, avg=average.

This classification confirmed the observed trends for gene expression, gene size, 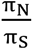, GC% and ENC (Figure 2A-E), and further revealed that core and dispensable genes differ markedly in their relative proportions of transcriptional categories (Figure 2F). Most core genes were VEGs (37.64%) or SEGs (33.49%), with only 28.87% classified as OOGs. By contrast, dispensable genes showed a comparable proportion of VEGs (33.39%) but a smaller proportion of SEGs (23.21%) and a higher proportion of OOGs (43.4%). Both gene sets contained representatives of all three expression classes, indicating that dispensable genes can adopt diverse transcriptional states, including stable expression.

**Figure 2:**
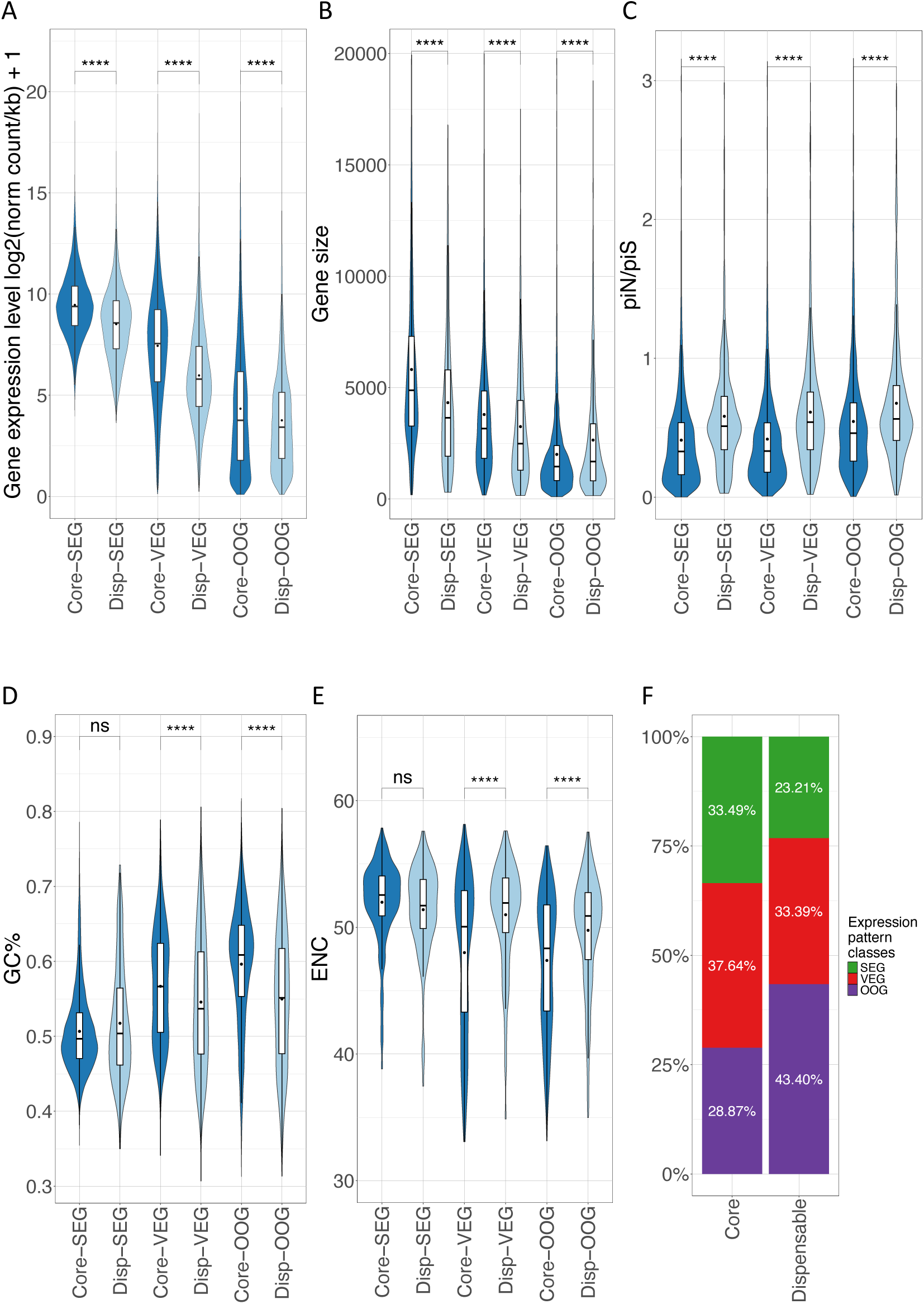
Features of core and dispensable by transcriptional categories. Violin plots for gene size (A), expression levels (B), 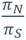 (C), Proportion of GC in coding sequence (D), ENC (F). (F) Proportion of stably expressed genes (SEGs, green), variable genes (VEGs, red) and on-off genes (OOGs, purple) among core and dispensable genes. All results were obtained on the B73 genotype. (A-E). Core gene distributions are in dark blue, and dispensable genes are in pale blue. (A-D) p-values were obtained using t-test: ns=not significant, **** indicates a p-value lower than 0.001.

Because *Helitrons* are known to mediate gene movement, we examined whether they overlap dispensable genes more frequently than core genes. Notably, dispensable genes overlapped *Helitrons* at 4.6 times the rate of core genes (44% *versus* 15%; Fisher OR = 4.6, p < 2.2e^-16^). Among dispensable genes, SEGs showed the highest *Helitron* overlap frequency, followed by OOGs and VEGs (Supplementary figure 4).

### Gene expression is a key property to distinguish core from dispensable genes

To gain deeper insights into the properties of core and dispensable genes, we re-examined the features described above within each transcriptional category (SEG, VEG and OOG). Within each transcriptional category, core/dispensable differences in gene expression level, gene size, and 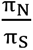 remained significant, but were smaller than the between-category differences observed within either gene set (Figure 2 A, B, D). A similar trend was observed for GC% and ENC, except that differences between core and dispensable genes were no longer significant within the SEG category (Figure 2 D, E). Furthermore, although dispensable genes displayed higher 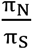 values than core genes on average, many dispensable genes featured 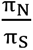 values below 0.5, indicating that they are likely subject to purifying selection. Similarly, a substantial subset of dispensable genes had ENC values comparable to those of core genes.

We therefore sought to identify the factors predictive of core or dispensable gene status. To this end, we first performed a principal component analysis. The first axis (PC1) which explained 38% of the variation, primarily distinguished transcriptional categories, with ENC and gene size being the most strongly contributive variables (Figure 3A). The second axis (PC2), which explained 28.6% of the variation, revealed a separation between core and dispensable genes, although their distributions largely overlapped (Figure 3B). This axis was primarily driven by the 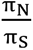 ratio and by expression level (Figure 3B).

**Figure 3:**
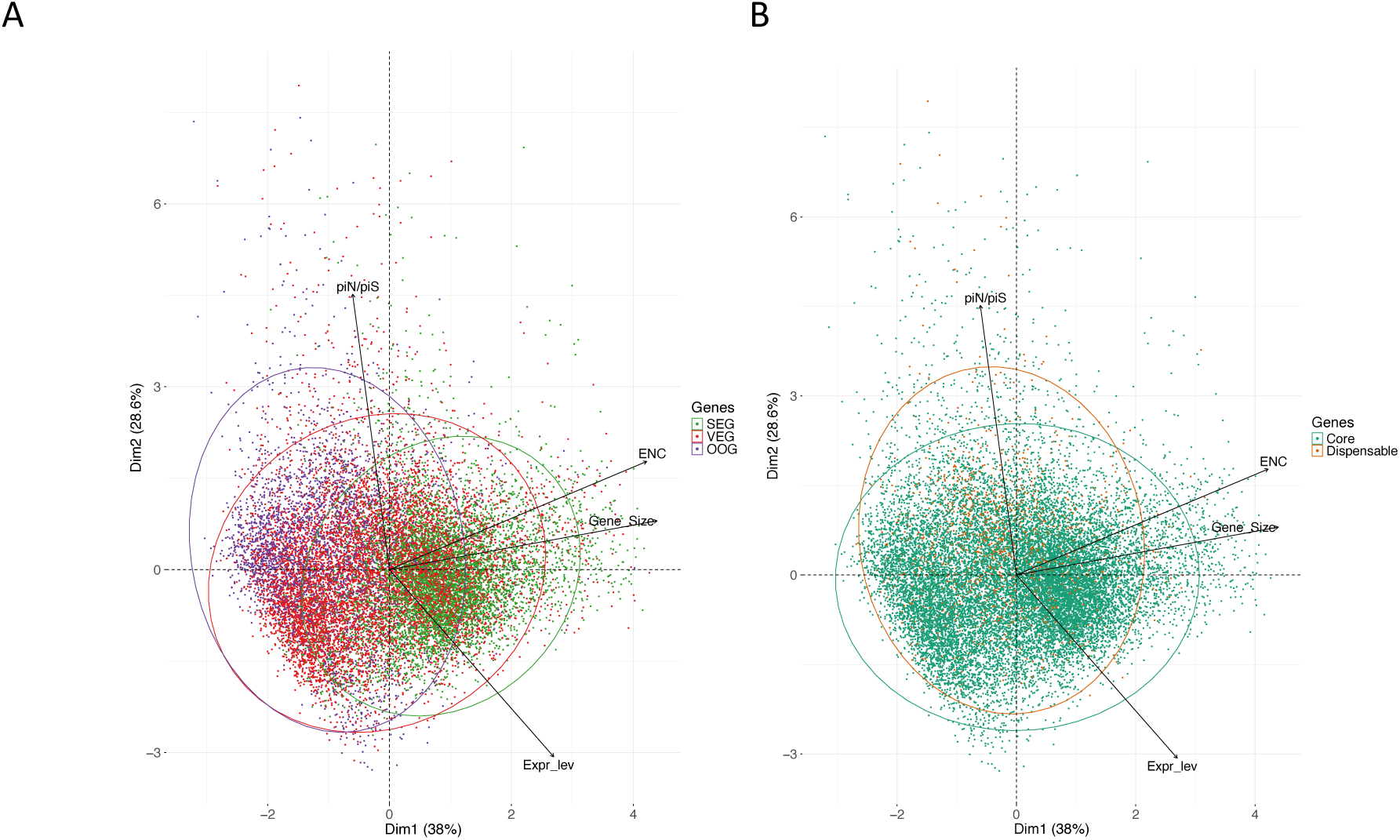
Principal component analysis computed from 4 variables on all gene. The PCA is computed on expression level,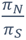, size and ENC. It is coloured by functional categories (A) and gene status and (B). Genotype is F2 and expression level were from well-watered tassel.

To assess the predictive power of the four variables, we fitted a logistic regression model (core/dispensable ∼ gene expression level + gene size +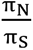+ ENC). All four variables contributed significantly to the models (Supplementary table 4); however, predictive performance was modest, with an average ROC-AUC of 0.69 (SD 0.2 over 141 glm, Supplementary table 5). This approach is nonetheless limited by potential misclassification of genes as core, given the inherently incomplete nature of a pangenome based on eight lines – genes currently classified as core may prove dispensable as additional genotypes are included – which may bias the training set and reduce predictive accuracy.

To further compare the relative contributions of the two main variables identified by the PCA – the 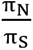 ratio and expression level – we sampled ‘dispensable-like’ genes from the core gene set based on each variable individually and on both combined. We then compared the distributions of these dispensable-like genes with those of actual dispensable genes across all four variables, treating the sampling variable as a control. Sampling based on either variable individually, or on both combined, partially shifted the distributions of dispensable-like genes towards those of dispensable genes, although differences persisted across all variables (Figure 4A-C). In contrast, when sampling was based on expression level alone or on both variables combined, the proportions of transcriptional categories in the dispensable-like set became indistinguishable from those of actual dispensable genes (Figure 4A and 4C). These findings are consistent with gene expression level being the primary factor distinguishing core from dispensable genes.

**Figure 4:**
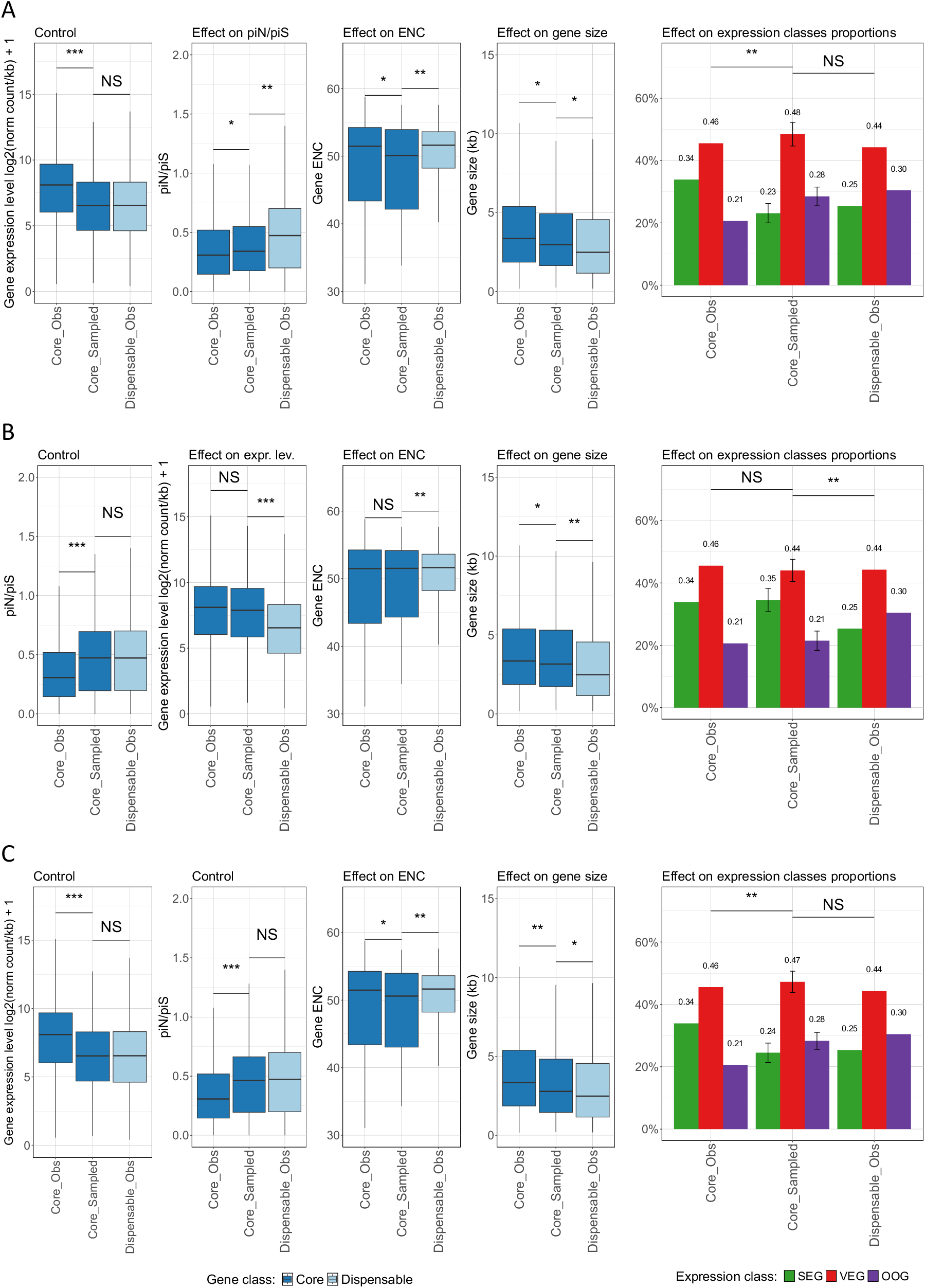
Effect of sampling dispensable-like genes from the core gene set on the distribution of gene features and relative proportions of transcriptional categories. Core genes are sampled according to dispensable genes expression level distribution (A), 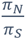 (B) and both gene expression level and 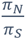 (C). p-values were obtained using t-test: ns=not significant, **** indicates a p-value lower than 0.001

### Dispensable and core genes share the same networks and functions

Because dispensable genes are, on average, shorter and less expressed than core genes, their functional annotation is known to be challenging. We were nevertheless able to assign at least one GO term to 44% of dispensable genes (compared with 51.5% of core genes), and could therefore test whether dispensable and core genes share biological functions using a GO term enrichment clustering approach. This analysis revealed a clear functional separation of transcriptional categories, indicating that our transcriptional classification captures meaningful biological distinctions (Figure 5). SEGs were predominantly associated with basal cellular functions (e.g., mRNA and protein metabolism, regulation, maintenance of subcellular components, and, to a lesser extent, energy homeostasis), VEGs were associated with environmental responses (e.g., photosynthesis regulation, response to karrikin and chitin), and OOGs were associated with cellular organisation, secondary metabolism and ovule protection, consistent with their tissue-specific expression.

**Figure 5:**
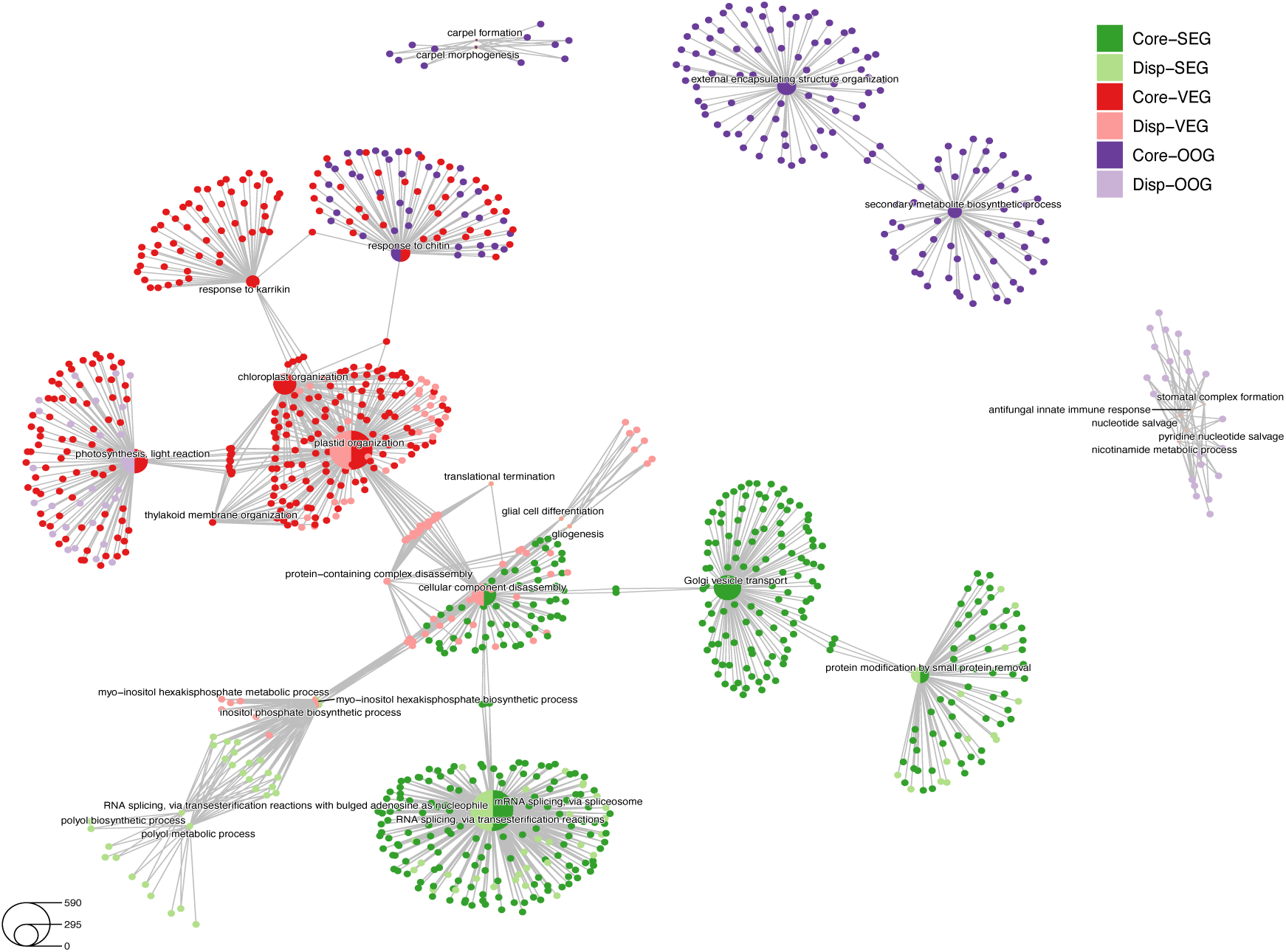
Cnet plot of over-represented GO terms (Biological processes) for cluster of genes grouped according to their genomic status and their expression pattern. Satellite dots are genes and are coloured following gene attributes (see legend), central pie charts are GO terms with proportion of core (dark) and dispensable (pale) genes linked to the term.

Core and dispensable genes frequently co-occurred in the same gene network clusters, and in similar proportions within those clusters. This was particularly evident in SEG clusters associated with mRNA splicing and protein modification, and in a VEG cluster involved in plastid organisation (Figure 5). The photosynthesis light-reaction cluster contained both core VEGs and dispensable OOGs, while the chitin-response cluster comprised exclusively core genes (both VEGs and OOGs). A small subset of functional clusters was composed entirely of dispensable genes, including those associated with inositol metabolism, translational termination (dispensable SEGs and VEGs), nucleotide salvage (dispensable OOGs) and neutral lipid/acylglycerol catabolism (dispensable VEGs). Taken together, these results demonstrate that, like core genes, dispensable genes participate in a broad range of biological functions, encompassing both basal cellular metabolism and environmental responses.

### Gene duplication occurs preferentially in stably-expressed genes and in dispensable genes

Having observed that a subset of dispensable genes is involved in basal cellular functions, we reasoned that their loss might be tolerated through functional redundancy, which could notably occur through the existence of paralogous copies. We therefore investigated the extent to which dispensable genes derive from gene duplication events, with particular interest in SEGs given their association with basal functions. To this end, we estimated the proportion of duplicated proteins in the pan-proteome inferred from our expressed pan-gene set using a protein clustering approach. Focusing on cases of substantial functional redundancy, two proteins were considered duplicates if they shared at least 80% sequence similarity over at least 60% of their length. Using these criteria, we identified 11,662 pan-proteins (31.45%) as being duplicated (Table 1). The proportion of duplicated proteins was significantly highest (χ^2^ test, p < 10^-123^) for SEGs (39.71%), followed by VEGs (29.92%) and OOGs (25.62%), indicating greater functional redundancy among SEGs, consistent with their involvement in basal biological functions (Table 2). Considering SEGs only, core genes appeared more often duplicated than dispensable genes (41.47% *versus* 35.39% respectively, Z-tests, p < 10^-8^ after Bonferroni correction, Table 3).

**Table 1:** Fraction of genes found in one or more copies according to their genomic classe (core or dispensable).

| pan-gene set (expressed) |  |  |  |
| --- | --- | --- | --- |
| Gene count | core | dispensable | total |
| single | 18,200 | 7,217 | 25,417 |
| duplicated | 7,349 | 4,313 | 11,662 |
| total | 25,549 | 11,530 | 37,079 |
| Fraction of core vs. dispensable genes (%) |  |  |  |
| single | 71.61 | 28.39 | 100.00 |
| duplicated | 63.02 | 36.98 | 100.00 |
| total | 68.90 | 31.10 | 100.00 |
| Fraction of single vs. duplicated genes (%) |  |  |  |
| single | 71.24 | 62.59 | 68.55 |
| duplicated | 28.76 | 37.41 | 31.45 |
| total | 100.00 | 100.00 | 100.00 |
| Fraction of each classe vs. total gene count (%) |  |  |  |
| single | 49.08 | 19.46 | 68.55 |
| duplicated | 19.82 | 11.63 | 31.45 |
| total | 68.90 | 31.10 | 100.00 |

**Table 2:** Fraction of genes found in one or more copies according to their expression pattern (SEG, VEG or OOG).

| pan-geneset (expressed) |  |  |  |  |
| --- | --- | --- | --- | --- |
| Gene count | SEG | VEG | OOG | Total |
| single | 6,772 | 9,438 | 9,207 | 25,417 |
| duplicated | 4,461 | 4,029 | 3,172 | 11,662 |
| total | 11,233 | 13,467 | 12,379 | 37,079 |
| Fraction of core vs. dispensable genes (%) |  |  |  |  |
| single | 26.64 | 37.13 | 36.22 | 100.00 |
| duplicated | 38.25 | 34.55 | 27.20 | 100.00 |
| total | 30.29 | 36.32 | 33.39 | 100.00 |
| Fraction of single vs. duplicated genes (%) |  |  |  |  |
| single | 60.29 | 70.08 | 74.38 | 68.55 |
| duplicated | 39.71 | 29.92 | 25.62 | 31.45 |
| total | 100.00 | 100.00 | 100.00 | 100.00 |
| Fraction of each classe vs. total gene count (%) |  |  |  |  |
| single | 18.26 | 25.45 | 24.83 | 68.55 |
| duplicated | 12.03 | 10.87 | 8.55 | 31.45 |
| total | 30.29 | 36.32 | 33.39 | 100.00 |

**Table 3:** Fraction of genes found in one or more copies according to their genomic classe (core or dispensable) and expression pattern (SEG, VEG or OOG).

|  |  | pan-gene set (expressed) |  |  |  |  |  |  |
| --- | --- | --- | --- | --- | --- | --- | --- | --- |
|  |  | SEG |  | VEG |  | OOG |  |  |
| Gene count |  | core | dispensable | core | dispensable | core | dispensable | total |
|  | single | 5,043 | 1,729 | 7,000 | 2,438 | 6,157 | 3,050 | 25,417 |
|  | duplicated | 3,514 | 947 | 2,617 | 1,412 | 1,218 | 1,954 | 11,662 |
|  | total | 8,557 | 2,676 | 9,617 | 3,850 | 7,375 | 5,004 | 37,079 |
| Fraction of core vs. dispensable genes (%) |  |  |  |  |  |  |  |  |
|  | single | 74.47 | 17.32 | 74.17 | 25.83 | 66.87 | 33.13 |  |
|  | duplicated | 87.77 | 23.50 | 64.95 | 35.05 | 38.40 | 61.60 |  |
|  | total | 76.18 | 19.87 | 71.41 | 28.59 | 59.58 | 40.42 |  |
| Fraction of single vs. duplicated genes (%) |  |  |  |  |  |  |  |  |
|  | single | 58.93 | 64.61 | 72.79 | 63.32 | 83.48 | 60.95 | 68.55 |
|  | duplicated | 41.47 | 35.39 | 27.21 | 36.68 | 16.52 | 39.05 | 31.45 |
|  | total | 100.00 | 100.00 | 100.00 | 100.00 | 100.00 | 100.00 | 100.00 |
| Fraction of each classe vs. total gene count (%) |  |  |  |  |  |  |  |  |
|  | single | 13.60 | 4.66 | 18.88 | 6.58 | 16.61 | 8.23 | 68.55 |
|  | duplicated | 9.48 | 2.55 | 7.06 | 3.81 | 3.28 | 5.27 | 31.45 |
|  | total | 13.85 | 4.33 | 15.57 | 6.23 | 11.94 | 8.10 | 100.00 |

## Discussion

To better understand the genomic, transcriptomic and functional features of maize dispensable genes, we constructed a pan-gene set from the genome assemblies of eight maize inbred lines representing European and American genetic groups. Predicted gene models ranged from 60,783 to 63,606 per genotype, substantially more than the average of 40,621 genes predicted for American NAM founder lines^23^, a difference attributable to the protein length filter applied in that study, which excluded genes encoding proteins shorter than 150 amino acids. We deliberately omitted this filter, as dispensable genes are on average shorter than core genes so that such a cutoff would systematically bias their analysis. Consistently, our dataset includes on average ∼20,000 coding genes shorter than 150 amino acids per genotype, 62% of which were present across all eight genotypes. With 76,358 unique gene models, our pan-gene set is consistent with previous estimates for maize, including those derived from transcriptome assemblies of 500 individuals (∼63,000-103,033 genes)^24^ and from whole genome assemblies comparison of 26 maize lines^23^, thereby confirming established estimates of the maize pan-gene repertoire, and highlighting the excellent sampling choice of our 8 lines.

Dispensable genes overlapped *Helitrons* at 4.6 times the rate of core genes (44% *versus* 15%). Although *Helitrons* prediction is notoriously challenging owing to the absence of strong terminal sequence signatures, one study reported that 90% of predictions made with the EDTA software were accurate^25^. *Helitrons* make up at least 2% of the maize genome and frequently carry gene fragments from hundreds of different genes^26^. By combining captured fragments, they can promote exon shuffling and the generation of novel transcripts, potentially giving rise to new proteins and contributing to genetic diversity^27^. *Helitrons* preferentially accumulate near gene-rich regions but avoid inserting directly into genes, and their gene fragment captures can be under purifying or adaptive selection, suggesting functional importance^27^. Our results indicate that a substantial fraction of dispensable genes originates from *Helitron*-mediated gene capture. Previous studies have reported that *Helitron*-captured genes can be highly expressed and show tissue-specific patterns^28,29^. Consistent with this, *Helitron*-overlapping dispensable genes were represented across all three transcriptional categories (SEG, VEG and OOG), although SEGs were moderately over-represented among them.

As repeatedly observed in several species, dispensable genes differ from core genes across several molecular and evolutionary features, exhibiting lower average expression levels and smaller gene size, alongside higher 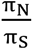 and ENC values. The distributions of these features however overlap substantially between the two gene sets, thus making it difficult to assign core or dispensable status to any individual gene on the basis of a single feature. Our analyses nevertheless suggest that expression level and to some extent 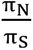 are the main factor distinguishing core from dispensable genes.

A notable finding is that all three transcriptional categories that we set up – SEG, VEG and OOG – contain dispensable genes, demonstrating that dispensable genes, like core genes, span the full range of expression patterns. Core and dispensable genes do, however, differ markedly in the relative proportions of these categories: dispensable genes are enriched in OOGs and depleted in SEGs compared with core genes. When transcriptional category is taken into account, expression level differences between core and dispensable genes persist but are substantially reduced relative to those observed without stratification. Furthermore, expression level differences are greater between transcriptional categories than between core and dispensable gene sets within any given category, with OOGs being the least expressed, followed by VEGs and SEGs. Altogether, these results indicate that the relative proportion in OOG or VEG genes together with the lower expression of these classes largely explains the overall tendency for dispensable genes to be expressed at lower levels than core genes. Taken together, these results suggest that the well-documented tendency for dispensable genes to be expressed at lower levels than core genes is largely a consequence of their higher proportion of OOGs and VEGs – inherently less-expressed gene classes – rather than a universal reduction in expression level *per se*. We therefore propose that examining the transcriptional category composition of core and dispensable genes may be more informative for understanding their biological functions and evolutionary dynamics than relying on expression level alone. Finally, the higher proportion of OOGs and VEGs among dispensable genes is consistent with potential roles in developmental or environmental responses. Yet, the observation that approximately one quarter of dispensable genes are SEGs indicates that a non-negligible fraction is involved in more basal biological processes than has previously been assumed.

GO-based functional annotation is often incomplete and must be interpreted with caution, a limitation that is particularly acute for dispensable genes given their shorter length and lower sequence conservation. To address this challenge, we leveraged our transcriptional classification to group genes into functional clusters defined jointly by core/dispensable status, GO annotation and transcriptional category. This enabled a more refined characterisation of the biological processes in which each gene class participates. Strikingly, most clusters comprised genes from a single expression category, confirming that our transcriptional categories capture meaningful functional distinctions. Among clusters composed exclusively of SEGs, two – associated with protein metabolism and RNA splicing, respectively – contained core and dispensable genes in approximately equal proportions. Two additional clusters, composed entirely of dispensable SEGs, were associated with polyol metabolism. Polyols are osmolytes derived from photosynthesis that serve as long-distance carriers of carbon and energy in plants, contributing to reserve partitioning between source and sink tissues^30^, carbohydrate storage and coenzyme regulation^30,31^. These findings provide clear evidence for the involvement of dispensable genes in fundamental plant biological processes.

The contribution of dispensable genes to basal cellular functions raises the question of functional complementation: how can the loss of genes underpinning essential processes be tolerated? Plant genomes are renowned for their propensity to undergo whole-genome and segmental duplications, and in maize approximately 70% of genes are estimated to have duplicated copies, with around 40% of duplication events having occurred after the sorghum/maize divergence^32^. We show that SEGs are duplicated most frequently (39%), compared with VEGs (29%) and OOGs (25%). Stratifying by core/dispensable status reveals that core SEGs have the highest duplication rate (41.07%), followed by dispensable genes across all expression classes (35%-39%), and then by core VEGs (27%) and core OOGs (16%). The elevated duplication rate among SEGs suggests that functional redundancy may be particularly important for safeguarding the expression of genes with basal roles. Within the SEG class, dispensable genes are slightly less duplicated than core genes (35.39% *versus* 41.07%), which is contrary to the expectation that the loss of a gene with a basal function would be buffered by the presence of a paralogous copy. Moreover, duplication rates among dispensable genes are similar across all three expression categories, suggesting that the relationship between dispensability and gene duplication is largely independent of gene function. Altogether, these results indicate that gene duplication only partially accounts for the dispensability of genes involved in basal functions, implying that other levels of biological redundancy must also contribute.

## Conclusion

Using a comprehensive pan-genomic and transcriptomic dataset from eight maize inbred lines, we investigated the genomic, evolutionary, and functional features of core and dispensable genes. Our results demonstrate that classifying genes into stably expressed, variably expressed, and on-off transcriptional categories is a powerful framework for capturing differences between core and dispensable genes, revealing that differences in class proportions largely account for the lower average expression observed in the dispensable gene set. Gene expression level and, to a lesser extent, relaxed purifying selection also contribute to distinguishing dispensable from core genes, therefore challenging the hypothesis that gene length could be a major determinant of dispensability.

*Helitrons* emerge as important drivers of dispensable genome dynamics, with dispensable genes overlapping *Helitrons* at 4.6 times the rate of core genes, implicating *Helitron*-mediated gene capture as a prevalent mechanism of dispensable gene formation. Functional network analysis further reveals that dispensable genes are not confined to environment-responsive or developmental roles: they participate in basal biological functions alongside core genes and within the same molecular networks. Protein clustering analysis suggest that only a fraction of dispensable gene functions are complemented through gene duplication, pointing to other levels of biological redundancy. Together, these findings advance our understanding of the forces shaping the dispensable genome and underscore the underappreciated functional relevance of genes classically considered non-essential.

## Material and methods

### Genetic material

Seven inbred maize lines (*Zea mays* ssp. mays) were selected to represent the main genetic American and European genetic groups. The panel comprises: the reference line B73, representing the Iowa Stiff Stalk Synthetic (BSSS) subgroup; MBS847, a dent line derived from the hybrid Pioneer3901 (Iodent subgroup); F252, an early-flowering dent line of Lancaster/Iodent origin; GF111, a Northern Flint line derived from the North American Gaspé population; F4, a Northern Flint line descended from the northern European population ‘Étoile de Normandie’; F2, a European Flint line originating from the southern French ‘Lacaune’ population; F331, a tropical line derived from high-altitude American germplasm; and EA1197, a tropical Spanish line of derived from the Southern Europe ‘Mollar de Almería’ population. A comprehensive description of each line, including genetic group, subgroup, pedigree, developer, and germplasm accession number, is provided in Supplementary table 1.

### Plant culture and transcriptomic data generation

Tissue samples were collected across six developmental stages: four days after sowing (4DAS), two-ligulated-leaf (V2), post-tassel initiation (PTI), pollen shed (PS), 12 days after pollination (12DAP), and 35 days after pollination (35DAP). Each genotype was represented by three biological replicates per treatment. For all sampling time points except 4DAS, one biological replicate corresponded to a single plant. For root and hypocotyl tissues collected at 4DAS, one replicate consisted of a pool of three plants.

Seedlings sampled at 4DAS were grown in the RGP greenhouse in 1-L perlite pots under half dark conditions for four days. Plants sampled at V2, PTI, and PS were grown at the PHENOARCH high-throughput phenotyping facility^33^, part of the Montpellier Plant Phenotyping (M3P), in 9-L polyvinyl chloride (PVC) pots (0.19 m diameter × 0.40 m height) filled with a 30:70 (v/v) clay-to-organic-compost mixture. Three seeds were sown per pot at a depth of 0.025 m and thinned to one plant per pot upon emergence of the third leaf. The greenhouse was maintained at 25 ± 3°C during the day and 18°C at night, under a 14-hour photoperiod. Supplemental lighting during day time was provided by 400-W HPS Plantastar lamps (OSRAM, Munich, Germany) at a density of 0.4 lamps m⁻² when natural solar radiation fell below 300 W m⁻², or to extend the photoperiod as required. Prior to silk emergence, ears were enclosed in paper bags to prevent uncontrolled pollination. PS-stage samples were collected under two contrasting water regimes. Well-watered (WW) plants were maintained at a soil water potential of −0.05 MPa. Water-deficit (WD) conditions were imposed after floral transition by progressively withholding irrigation until a soil water potential of −0.30 MPa was reached, which was thereafter maintained for the remainder of the experiment. Kernel samples were collected at 12DAP and 35DAP from field-grown plants at the INRAE experimental station of Saint-Martin-de-Hinx (France). A comprehensive overview of tissue types, growth conditions, and sampling design is provided in Supplementary Table 1. All samples were flash-frozen in liquid nitrogen immediately upon harvest and stored at −80°C.

Frozen tissues were ground in liquid nitrogen using a mortar and pestle. mRNA extraction, library preparation, sequencing, and raw data preprocessing were performed as previously described^34^.

### Transposable elements annotation

We first annotated the eight new genome sequences independently using the EDTA v1.7.3 pipeline (https://github.com/oushujun/EDTA). Briefly, we identified independently LTRs, TIRs and helitrons using the EDTA_raw.pl script, and then used the EDTA.pl script to merge the three consensus lists thus generated and a pre-existing library of maize consensus TE sequences (maizeTE02052020) available on the EDTA github, using the following options: --species Maize --anno 1 --overwrite 0 --threads 47 –curatedlib maizeTE02052020. We then merged the TE consensus sequences obtained for the eight genome sequences, and removed redundancy. To this end, we first identified potential duplicates by performing pairwise comparison of consensus sequences from all eight genomes. Sequences with divergence over 40 on at least 80% of their sequence, on at least 80 bp were considered unique. RepeatMasker allowed to identify consensus sequences with divergence over 40, using the following options: RepeatMasker -pa 24 -q -no_is -norna -nolow -div 40. We then kept only unique consensus sequences, using the cleanup_tandem.pl scripts from EDTA with the following options: - misschar N -nc 50000 -nr 0.8 -minlen 80 -minscore 3000 -trf 0 -cleanN 1 -cleanT 0. We considered as duplicates the pairs of TE consensus meeting the similarity criteria and either annotated with the same order and family ID, or annotated with a defined family and/or order for one member, but as unknown for the other. We also considered as duplicates DNA and MITE elements that met the similarity criteria. Finally, sequences that reached similarity criteria with several sequences from other or unknown families were considered as false positives and filtered out. For each set of duplicates, we retained only one consensus, using the alphabetical sorting of maize lines (first the TE consensus from the B73 library, then if absent, the one from the EA1197 library, etc.). This TE consensus list is available as a fasta file in supplementary data AMAIZING_maizeTEconsensus.fa.

Finally, we annotated each of the eight genome assemblies with this final consensus library using the EDTA_anno.pl pipeline with the following options: --overwrite 0 --species Maize --anno 1. TEs annotation for each of the eight genome is available on maizegdb.org (for instance, for line EA1197, https://download.maizegdb.org/Zm-EA1197-REFERENCE-AMZ-1.0/), and the code is available on the INRAE forge (see Data and code availability section).

### Gene prediction and functional annotation

Gene models were predicted independently for each genome assembly For each line, mRNA-seq reads were aligned to the corresponding genome assembly using Star^35^ (--sjdbOverhang 99). The resulting alignments were processed with StringTie^36^ v2.1.6 software (-f 0 -c 2.5 -g 50 –rf) to reconstruct a per-genotype transcript catalogue. Gene models were subsequently predicted with maker^37^ v4.3 software, using the genotype-specific transcript sequences described above as empirical evidence, supplemented by plant protein sequences retrieved from UniProt (release 2021_02). The MAKER pipeline integrated the following external tools: ncbi-blast-2.7.1+^38^, RepeatMasker^39^ 4.1.2, Exonerate^40^ 2.66.3, SNAP^41^ 2006-07-28, Augustus^42^ 3.0.1, tRNAscan-SE^43^ 2.0.1. Resulting gene models were filtered with the quality_filter.pl tool from the Maker 4.3 suite, retaining only models with an Annotation Edit Distance (AED) score below 0.5. Gene models entirely contained within predicted transposable element (TE) loci – with the exception of *Helitrons* –were further discarded, yielding a high-confidence gene model set in GFF format for each genome.

Protein sequences were extracted from the filtered GFF files using gffread^44^ v0.12.7. A non-redundant set of representative protein sequences was derived with CD-HIT^45^. These representative sequences were subsequently mapped onto each of the eight genome assemblies using miniprot^46^ 0.11, to recover gene models potentially missed during the initial prediction phase owing to a lack of RNA-seq read support in a given genotype. The full complement of representative protein sequences for each genome was then processed with to InterProScan 5^47^ v5.52-86.0 and eggNOG-mapper^48^ v2.1.7 to assign functional annotations. Gene models associated with Pfam domain identifiers characteristic of transposable elements were removed to obtain the final high-confidence gene model set for each genome. The corresponding GFF files were supplemented with functional annotation attributes. Codon usage statistics, including CDS GC content and the effective number of codons (ENC), were computed using the Cubar^49^ R package. The AED score distribution of the final gene models was homogeneous across genotypes and consistent with that obtained for the B73 NAM5 genome annotation (Zm00001eb.1) (Supplementary figure 5).

### Pan-gene set generation

Gene sets predicted for each genotype were compared to establish gene homology relationships across the eight genotypes and to derive a binary presence/absence matrix for all genes in the panel. Two complementary strategies were employed in parallel: one gene-centric approach aimed at identifying homologous gene pairs, and one sequence-based approach designed to detect homologous genomic regions. For the gene-centric approach, GeneSpace^50^ was used to infer homologous and paralogous gene relationships by integrating synteny information – based on the chromosomal positions of genes within orthologous groups (OGs) – as a constraint on the OG inference performed by OrthoFinder^51^. For the sequence-based approach, collinear genomic regions, hereafter referred to as anchor sequences, were identified using SibeliaZ^52^. Anchor sequences were required to satisfy two criteria: uniqueness within each genome assembly, and presence in at least one additional genotype. A total of 9,641,104 such anchor sequences were retained.

Results from both methods were combined to compute a set of pan-genes. Briefly, for each gene identified by GeneSpace in a given genotype, overlapping anchor sequences were used to search for homologous genes across the remaining genotypes, using in-house scripts. Genes for which homology assignments could not be resolved, or for which the two methods yielded discordant results, were excluded from the final pan-gene set (on average, approximately 2,000 genes per genotype).

### Gene expression analysis

For each sample, mRNA-seq reads were aligned to the genome assembly of the corresponding genotype using STAR^35^. STAR genome indices were built with the following parameters: --runMode genomeGenerate, --sjdbGTFfile (GTF-formatted gene annotation of the corresponding genome), --sjdbOverhang 99. Read alignment was performed with the following settings: --outSAMtype BAM SortedByCoordinate, --outSAMprimaryFlag OneBestScore, --alignIntronMin 5, --alignIntronMax 60000, --outFilterMultimapScoreRange 0, --outFilterMultimapNmax 20, --alignEndsType Local, --sjdbGTFtagExonParentGene gene_id. Per-gene read counts were obtained with featureCounts featureCounts^53^ v2.0.3 using the following options: -p -O -M --primary -s 2 -t exon -g gene_id.

To quantify gene expression levels, genes with absent or consistently low raw counts across all samples were first removed using HTSFilter^54^. Retained raw counts were normalised using the trimmed mean of M-values (TMM) method implemented in the edgeR^55^, and normalised counts were subsequently log₂-transformed as log₂(normalised count + 1).

For each genotype, expressed genes were further grouped into three classes according to their expression pattern across all samples of this genotype. Genes with a coefficient of variation (CV) of normalised expression across all samples below 0.1 were designated as Stably Expressed Genes (SEGs). Genes not detected in at least one sample were designated as On–Off Genes (OOGs). All remaining genes were classified as Variably Expressed Genes (VEGs).

### Gene ontology and functional analyses

A custom gene annotation database (org.Zmays.eg.db) was constructed using the AnnotationForge^56^ R package, incorporating Gene Ontology (GO) terms derived from the pan-gene set functional annotation described above. Genes were partitioned into six classes defined by the cross-classification of genomic status (core or dispensable) and expression category (SEG, VEG, or OOG, as defined in the previous section). These six gene lists, together with the annotation database, were supplied as input to the compareCluster function of the clusterProfiler^57^ R package, which identified enriched GO terms for each class using the enrichGO method. P-values were corrected for multiple testing using the Benjamini–Hochberg procedure, with a false discovery rate threshold of q < 0.05.

The output of the compareCluster analysis was then passed to the cnetplot function of the clusterProfiler package to generate concept–gene network graphs. In these networks, each node represents either an enriched GO term or an individual gene; edges connect a gene node to each GO term with which it is associated. Node size is proportional to the number of genes associated with the corresponding GO term, and gene nodes are coloured according to their expression category membership (figure 4). This representation simultaneously captures the associations between gene class membership – defined by expression category and core/dispensable status – and functional similarity among enriched GO terms, as quantified by semantic similarity scores.

### Principal component analysis

Principal component analysis (PCA) was performed on four genomic and transcriptomic variables characterised across the pan-gene set: normalised gene expression level, the 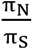, CDS size, and ENC. All variables were scaled to unit variance prior to analysis (scale.unit = TRUE), and the first six principal components were retained (ncp = 6), using the FactoMineR^58^ R package. Because gene expression levels vary across developmental stages and conditions, a separate PCA was computed for each sample independently. All per-sample PCAs yielded highly consistent results (data not shown); the PCA computed from a representative sample is presented in Figure 3. Genes were projected onto the principal component space and coloured according to either their pan-genomic status (core or dispensable) or their expression category (SEG, VEG, or OOG).

### Generalized linear models

The pan-genomic status (core or dispensable) of each gene was modelled as a binary response variable using a weighted generalised linear model (GLM) with a binomial error distribution, implemented via the glm() function in R. The four predictor variables were normalised gene expression level, CDS size, 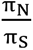, and ENC, yielding the following model formula:

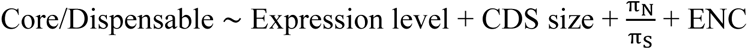

To account for the class imbalance between core and dispensable genes, a range of observation weights (18 to 26) was systematically evaluated, and the weight value yielding the best model performance was retained. A separate model was fitted for every combination of genotype, developmental sample, and growth condition available in this study. For each model, a stratified training set comprising 80% of the observations was constructed using the createDataPartition() function from the caret R package, with the remaining 20% reserved as a held-out test set. Model fit was assessed using the Akaike Information Criterion (AIC), computed with the AIC() function in R. Predictive performance was evaluated on the test set by computing the area under the receiver operating characteristic curve (AUC-ROC) using the auc() function from the pROC^59^ R package.

### Resampling of core genes to match dispensable gene parameter distributions

Core genes were sampled to reproduce the distribution of several parameters (ENC, CDS size, Gene 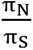, gene expression level) observed in dispensable genes. For each of the four parameters under study – ENC, CDS size, 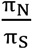, and normalised gene expression level –a subset of core genes was drawn without replacement (n = 300) using the sample() function in base R, such that the distribution of the focal parameter within the resampled subset matched that observed in dispensable genes. This procedure yielded four independent resampled core gene sets, each constrained to mirror the dispensable gene distribution for one parameter at a time.

For each resampled set, the values of all four parameters were computed. The distribution of the parameter used as the matching criterion was examined to verify the adequacy of the resampling. The distributions of the remaining three parameters within the resampled core gene subset were then compared to those of both the full (non-resampled) core gene set and the dispensable gene set using two-sample t-tests, to determine whether observed differences between core and dispensable genes persisted after controlling for the matched parameter.

### Gene molecular evolution

The 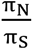 ratio was computed for all genes present in at least four genotypes and exhibiting no copy number variation. Protein sequences translated from the corresponding CDS were first aligned using MAFFT v7.505, called via the ips R package v0.0.11^60^ with the following settings: method = "auto", maxiterate = 0, op = 1.53, ep = 0. The underlying CDS sequences were then reverse-aligned against the protein alignment using the reverse.align() function from the seqinr R package v4.2.23, to obtain codon-partitioned nucleotide alignments. Pairwise πN and πS values were computed for all sequence pairs within each orthologous gene group using the kaks() function from the seqinr package, and the resulting pairwise estimates were averaged across all pairs within the group. A 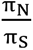 ratio was retained for each gene for which the mean pairwise πS exceeded zero, to exclude genes for which the ratio is undefined.

### Analysis of gene duplication rate

To estimate gene copy number within each genotype and across the pan-gene set as a whole, protein sequence sets were clustered using MMseqs2^61^ (v14-7e284) with the following settings: cluster, --cov-mode 0, -c 0.6, --min-seq-id 0.8. These parameters define a clustering threshold requiring at least 60% bidirectional sequence coverage and 80% sequence identity, enabling the grouping of gene copies likely arising from duplication events.

## Availability of data and materials

The sequencing reads and the genomes assembled from these data have been uploaded to the European Nucleotide Archive (ENA) at www.ebi.ac.uk/ena as part of the Amaizing project “De novo assembly and annotation of 7 maize inbred lines of importance for breeding programs in Central Europe”, and are accessible under project PRJEB89017.

Assembled genome sequences, as well as gene and TE annotations are available at https://www.maizegdb.org/amaizing_project. Transcriptomic data are available upon request to Clémentine Vitte and Johann Joets, and will be made publicly available upon article acceptance.

## Authors contribution

JJ designed the study. CV coordinated the project. JJ, CV and PR secured funding. JJ, CV, MF, MM and KB analysed the data. JJ and CV wrote the first draft of the manuscript. MF and MT discussed results and revised the manuscript. OT and CW coordinated water deficit experiments. LCB and OT conceived and performed PHENOARCH experiments. OT and SCo conceived plant sample collection procedure. CV and JJ collected plant samples for genome assembly. WM generated high quality DNA samples for genome assembly. OT, CV, JJ, AV, SCh, PR and CP collected samples for transcriptomic data generation. AR, AV, HB, SCo and JL generated mRNA samples for transcriptomic data generation. SCo, CV and SCh designed and assembled mRNA plate organization. SP generated transcriptomic data. CPLR assessed transcriptomic data quality, MLM and KB made preliminary data exploration analyses. VB deposited transcriptomic data to public databases.

## Acknowledgements

We thank Alain Charcosset for coordinating the Amaizing program and François Tardieu for advices and providing services of PHENOARCH facility. We are grateful to Cyril Bauland for expertise in maize germplasm accession nomenclature. We thank French maize inbred lines seed bank (CRB, INRAE Saint Martin de Hinx), Adrienne Ressayre and Christine Dillmann (GQE-Le Moulon) for providing seeds of F252 and MBS847, and Silvio Salvi (University of Bologna) for early access to seeds from the GF111 inbred line. We thank the Genotoul bioinformatics platform Toulouse Midi-Pyrenees for providing computing and storage resources, and Shujun Ou for providing an upgraded TE maize database that we used as starting point for TE annotation.

## Declarations

### Ethics approval and consent to participate

Not applicable

### Consent for publication

Not applicable

