## Supplementary information for "How to be dispensable: genomic and transcriptomic determinants in maize genes"

### Supplementary figures

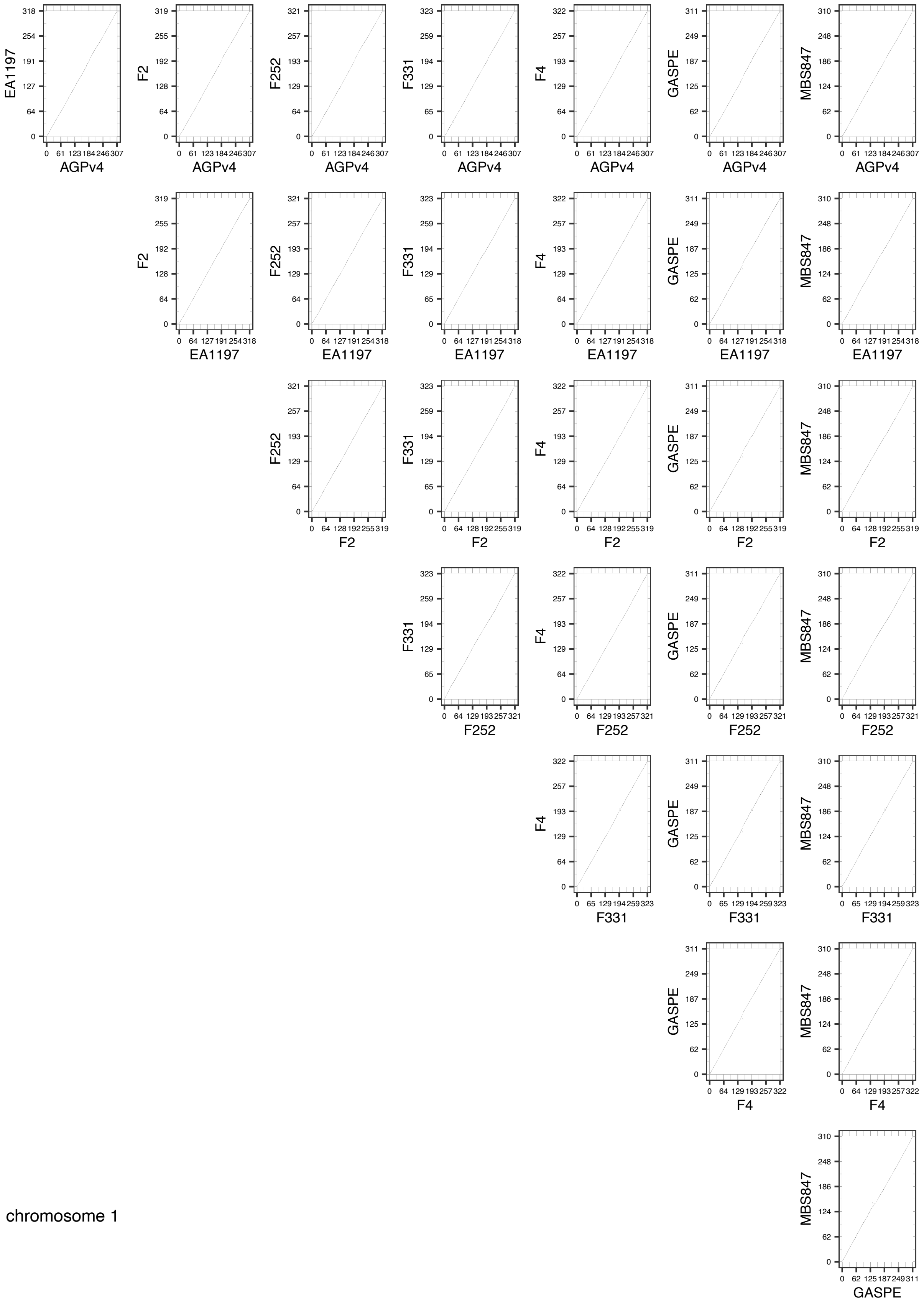

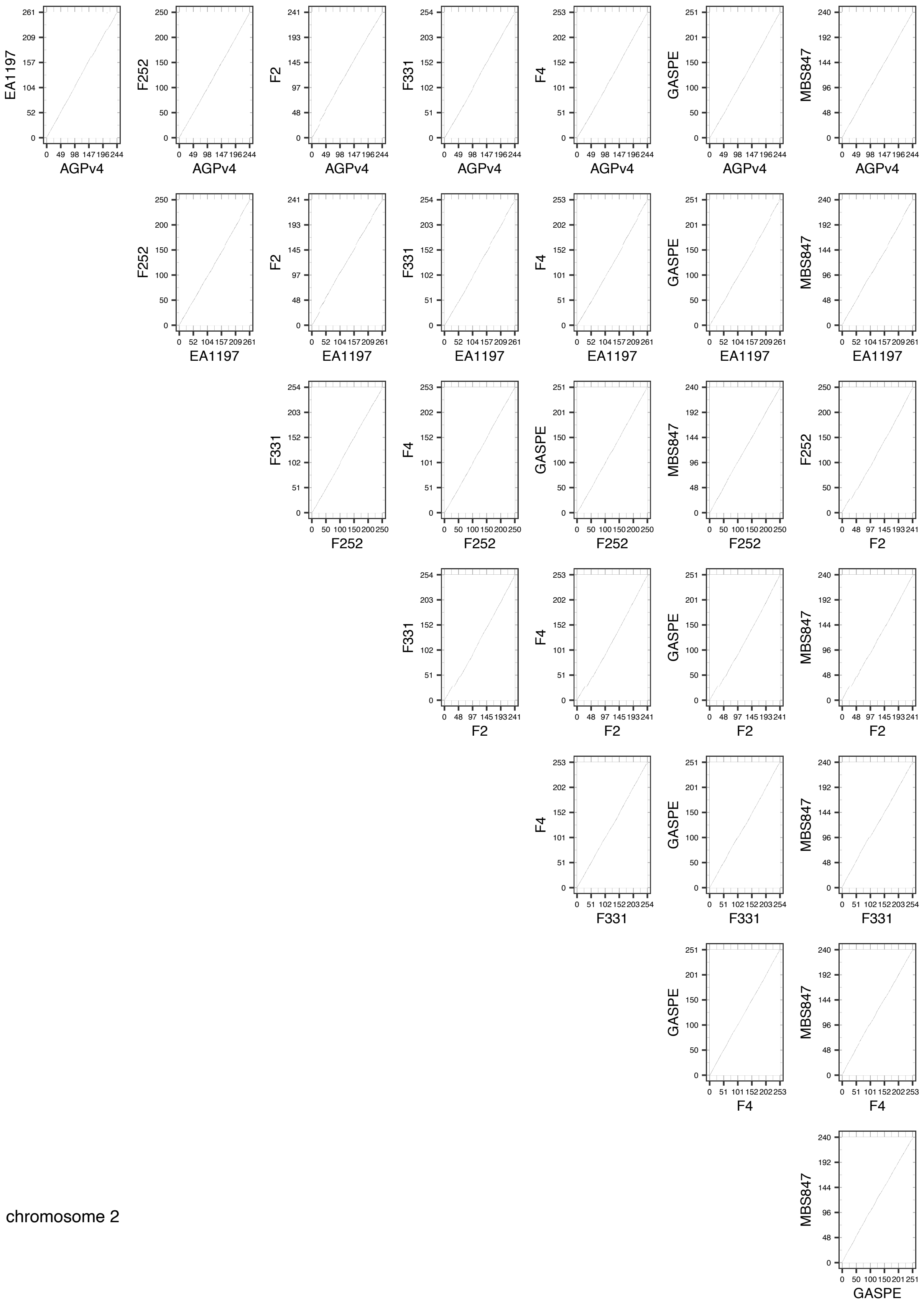


chromosome 2

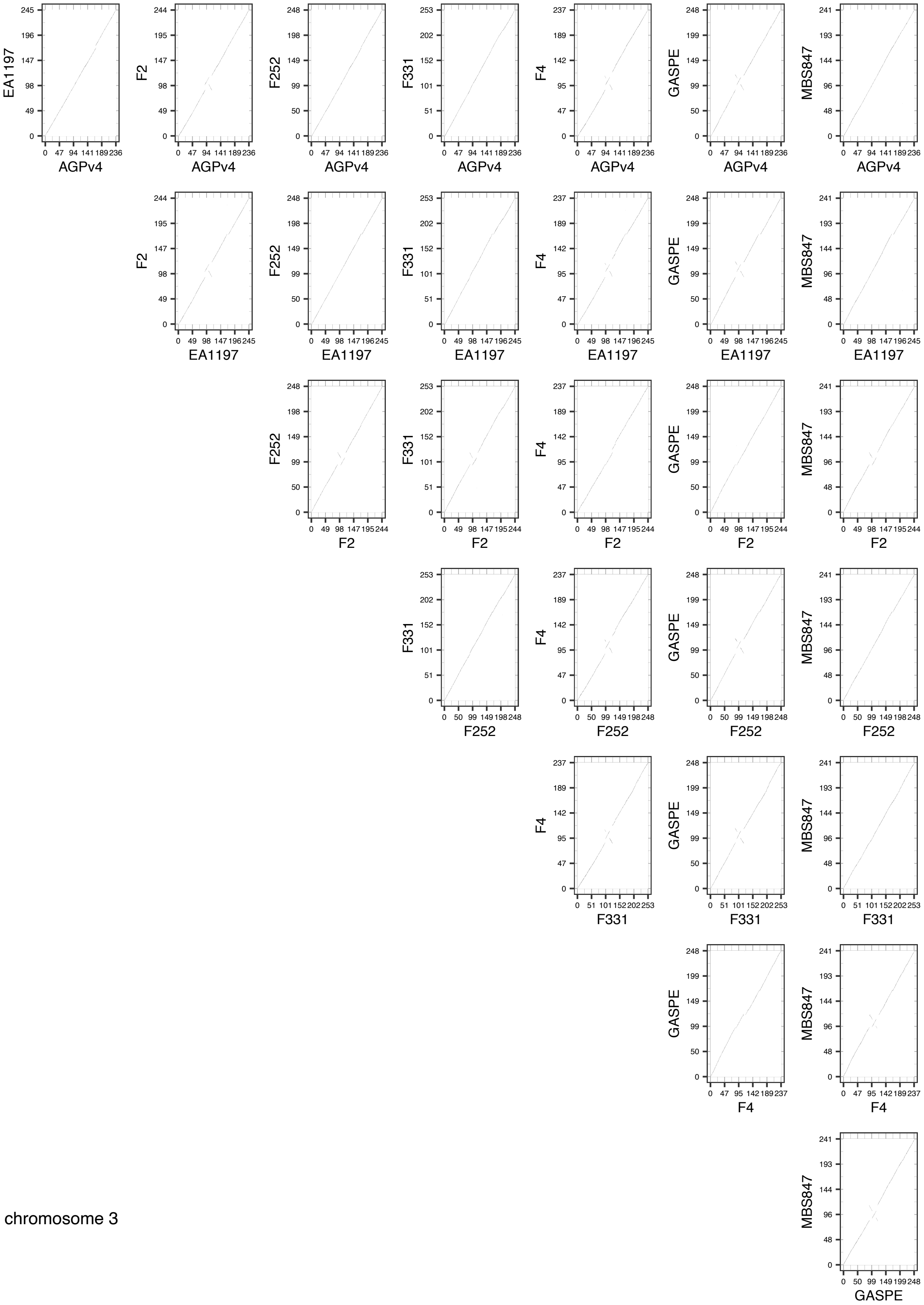


chromosome 3

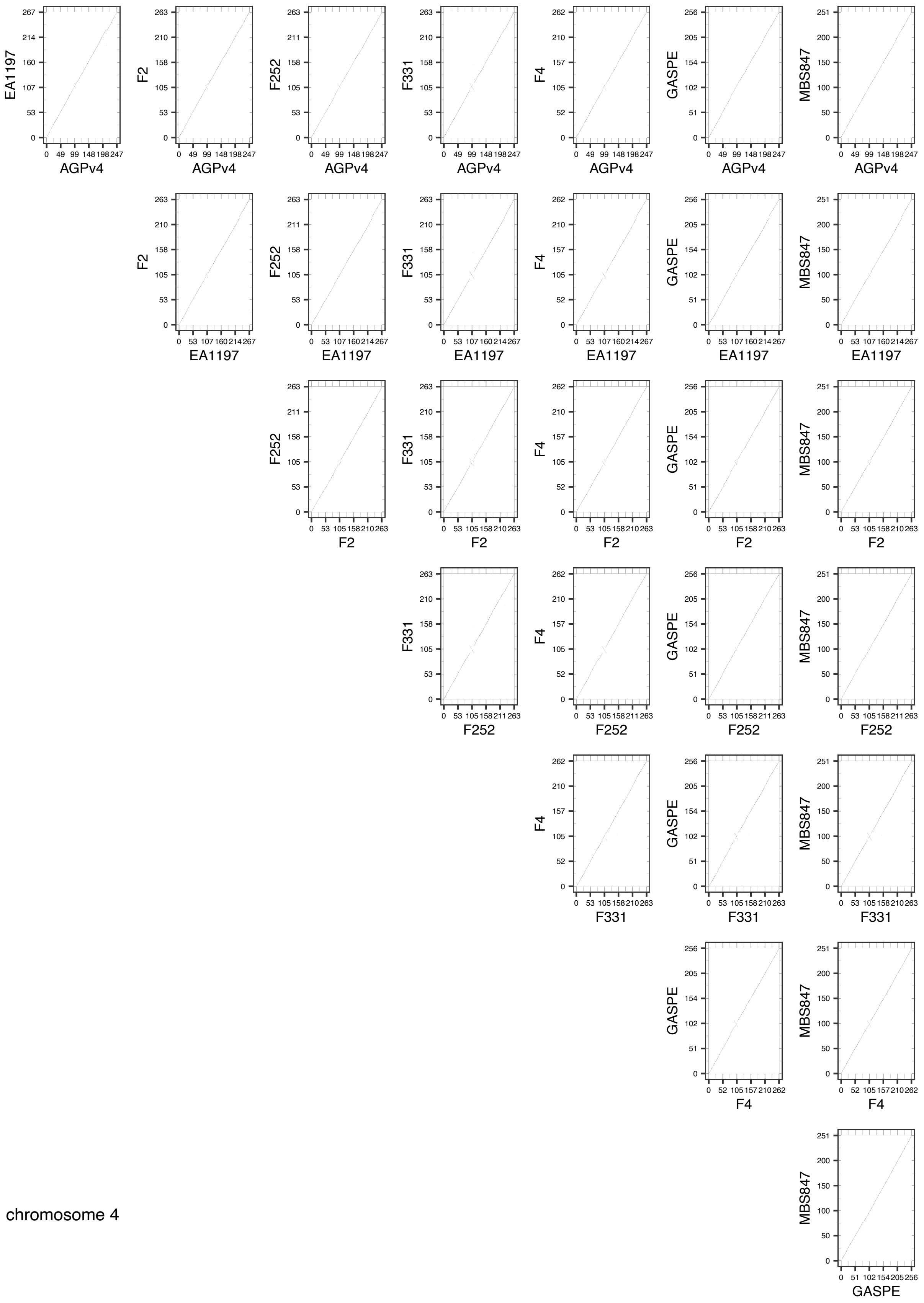


chromosome 4

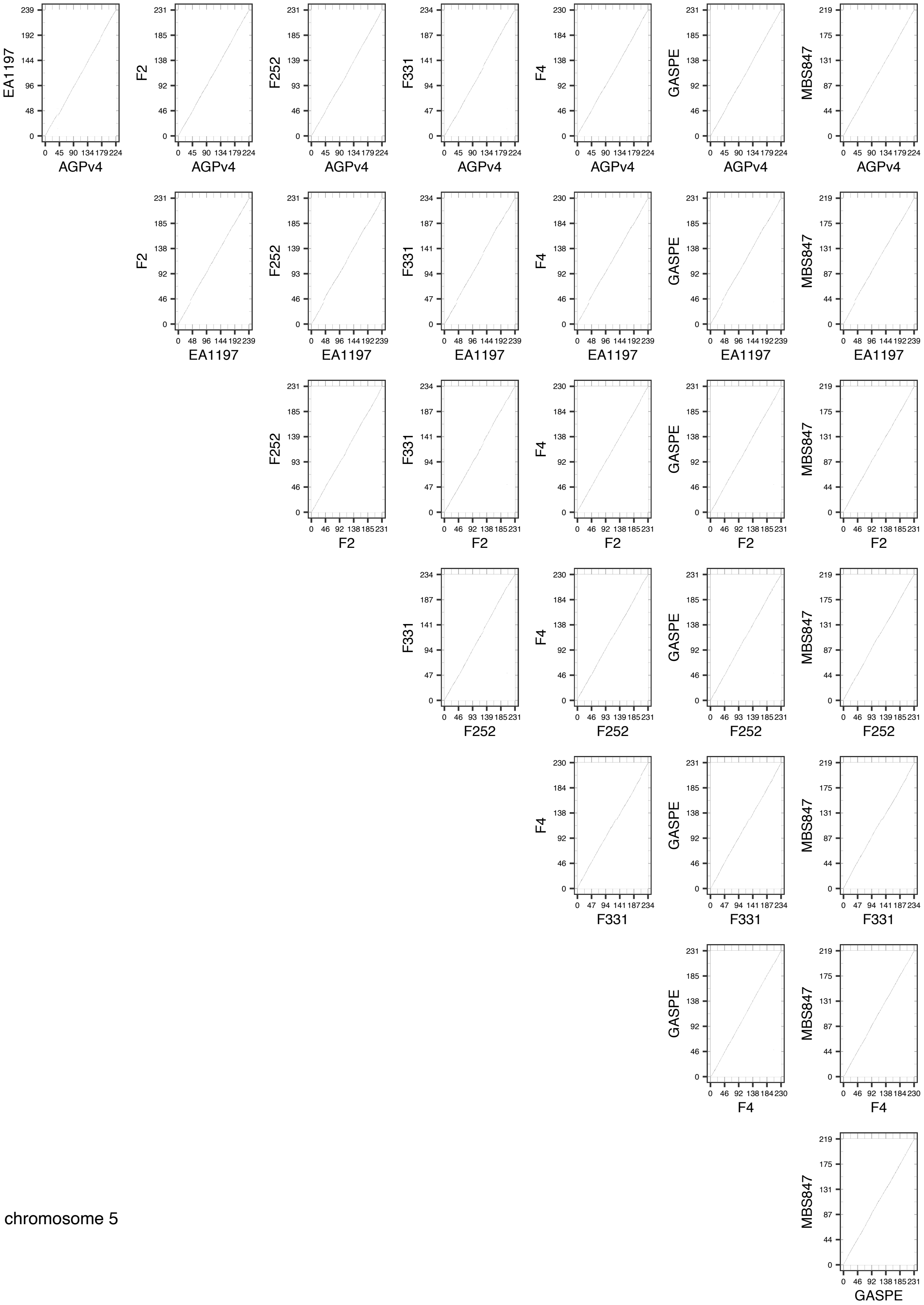


chromosome 5

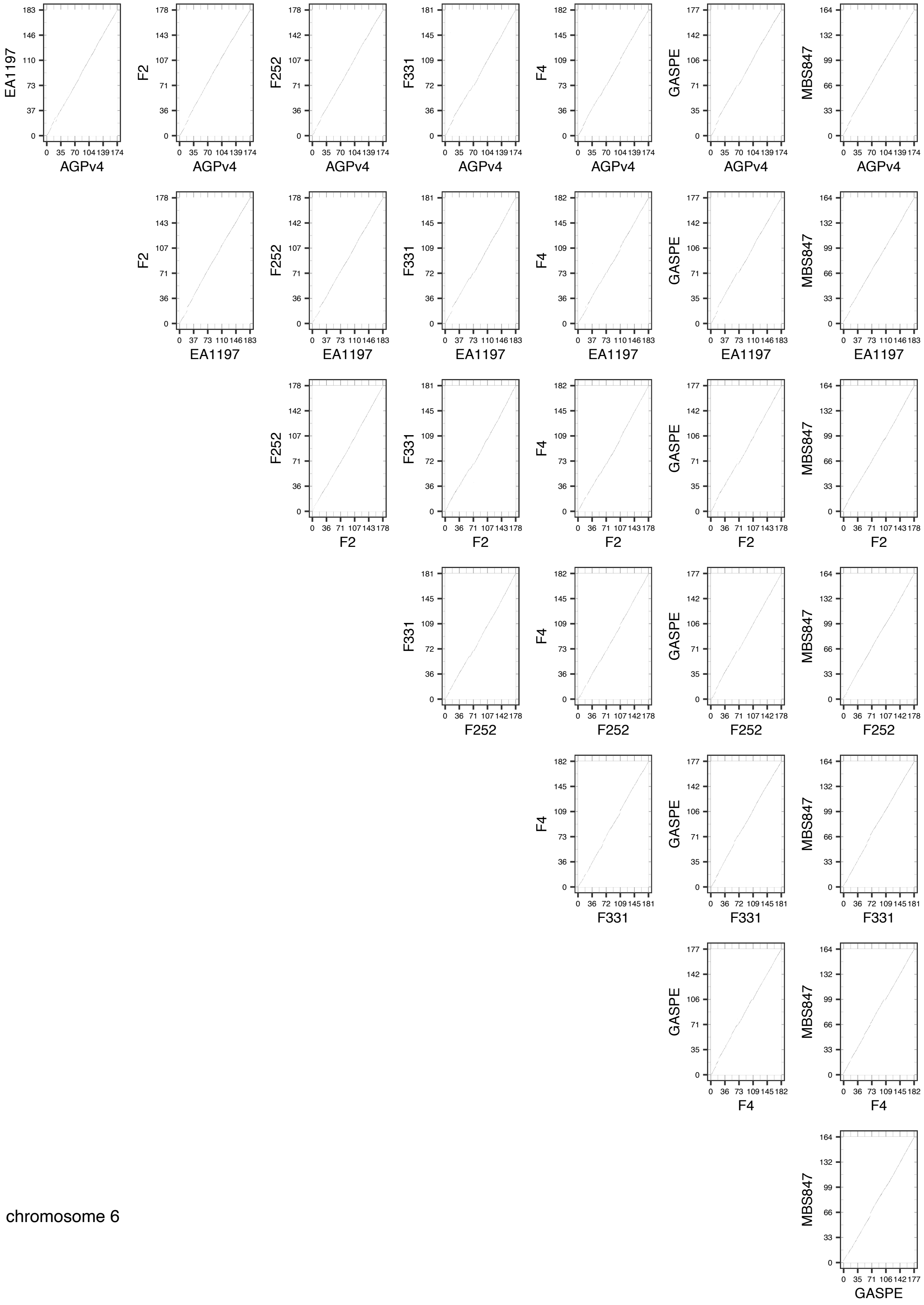


chromosome 6

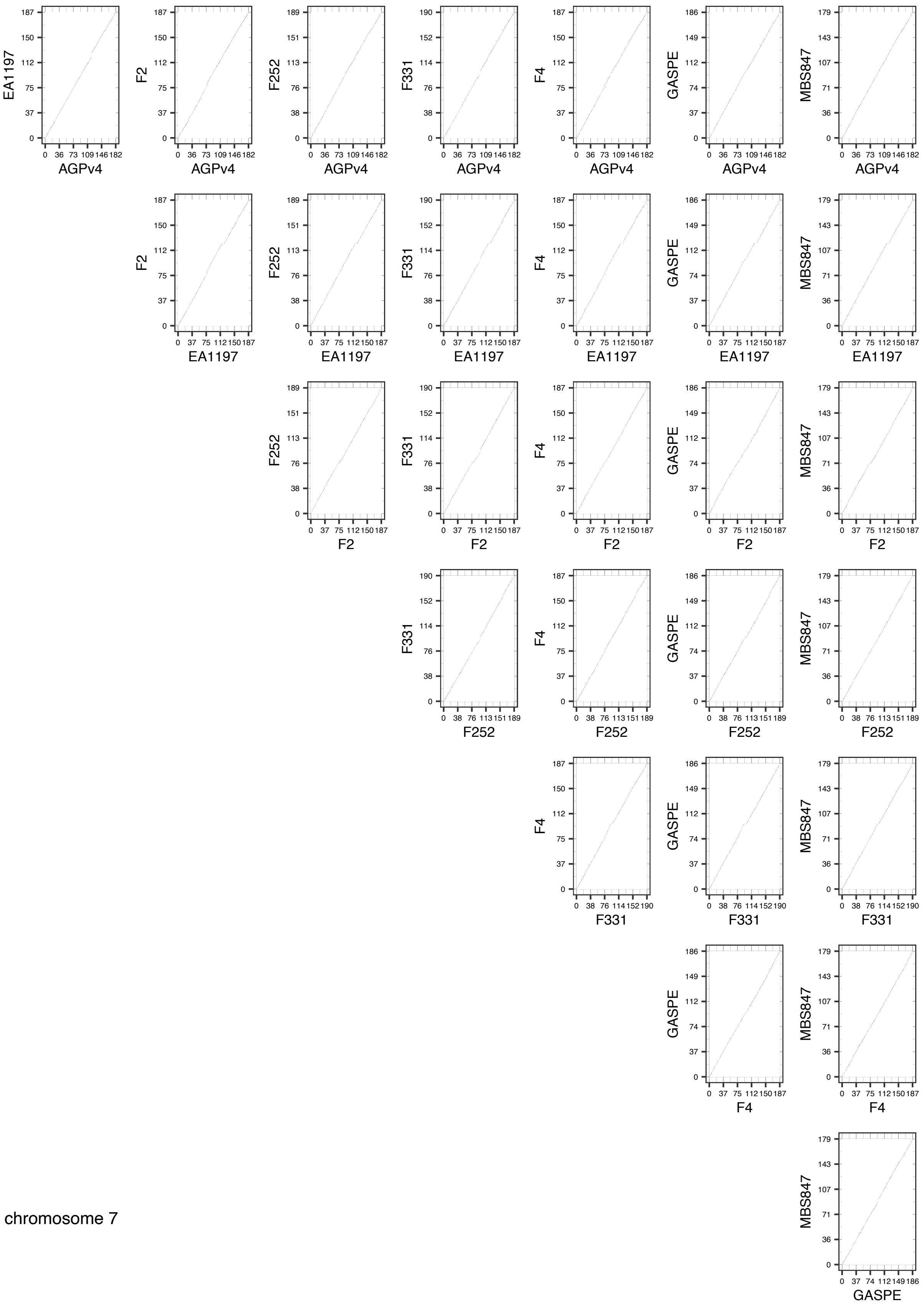


chromosome 7

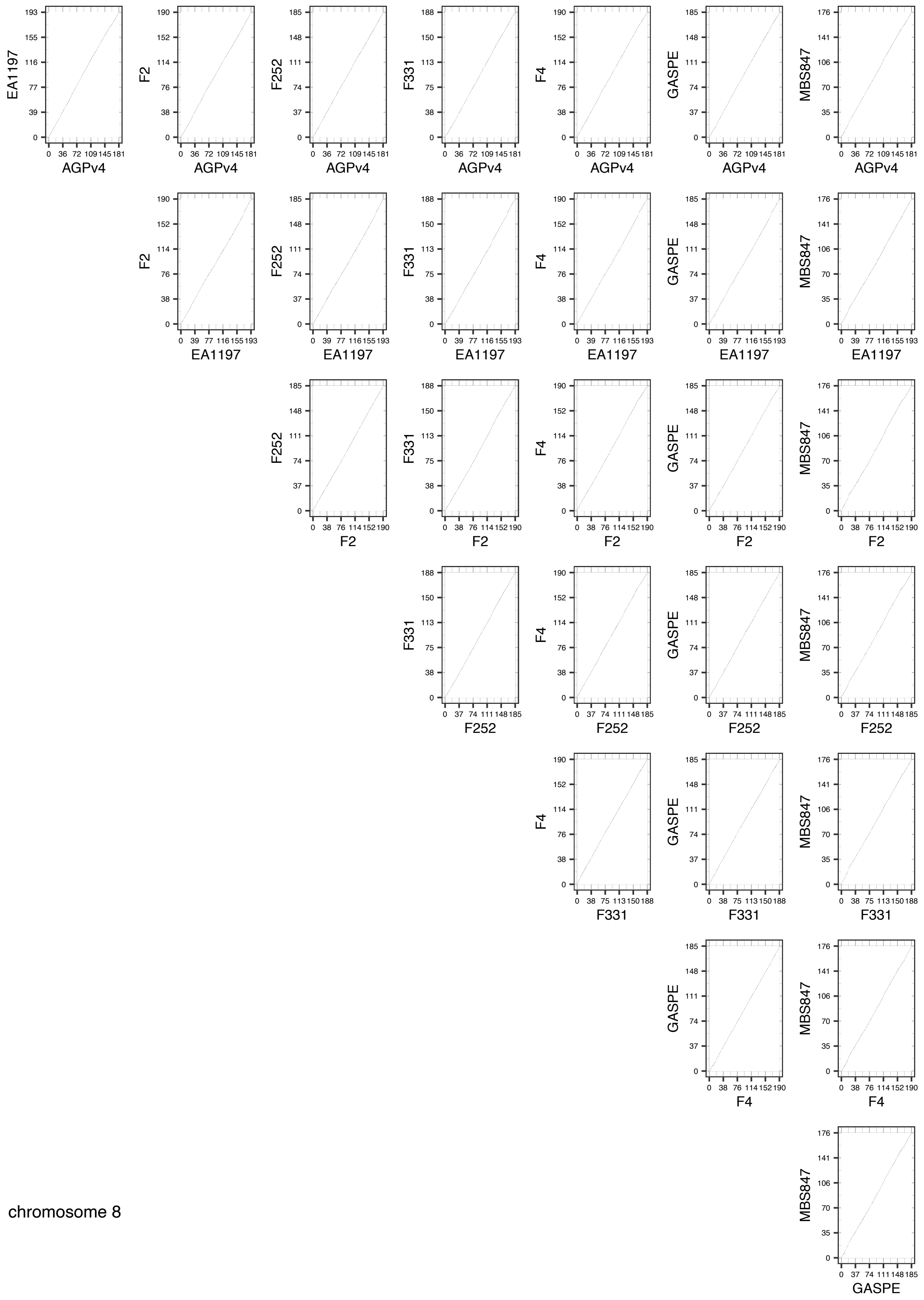


chromosome 8

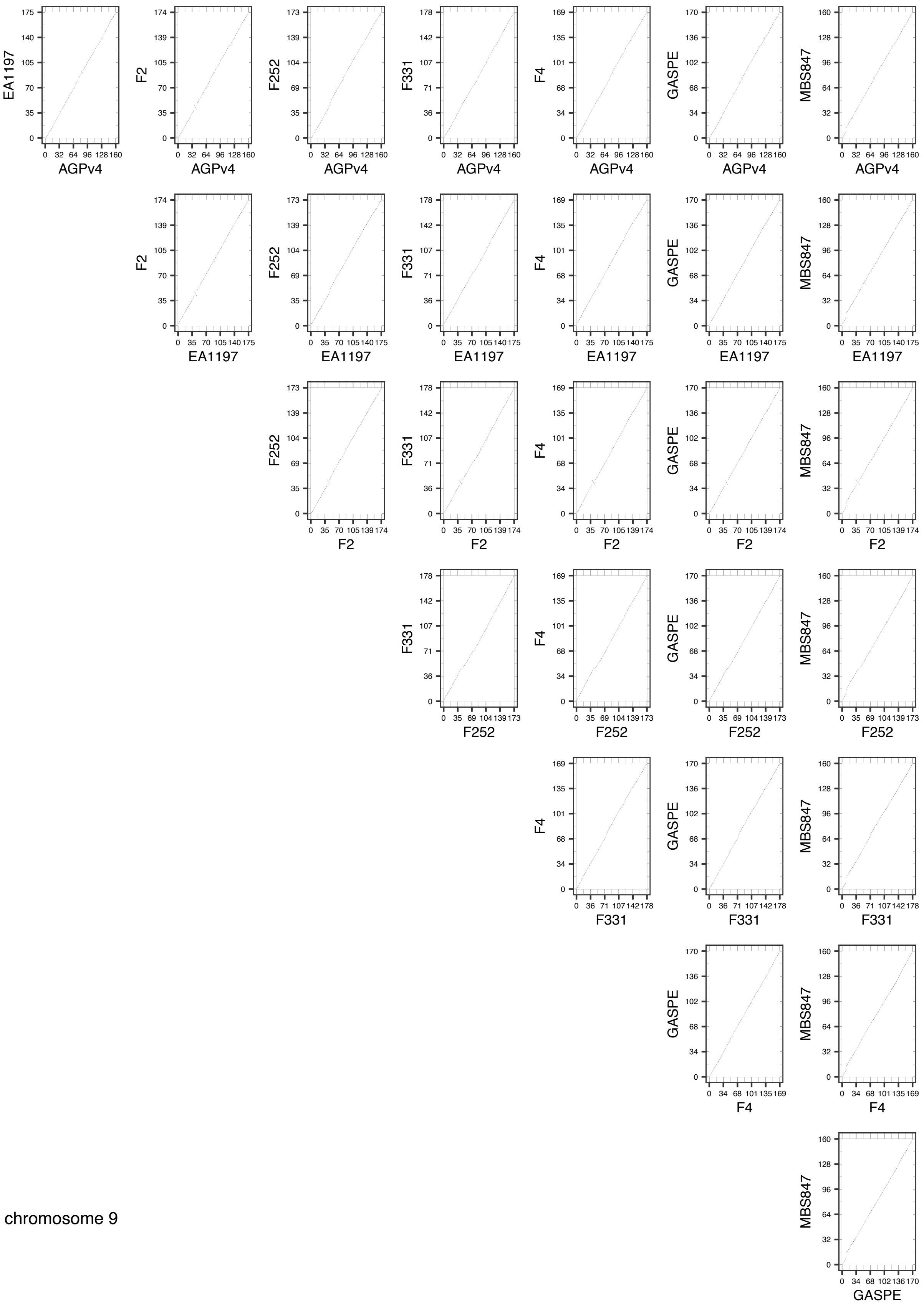


chromosome 9

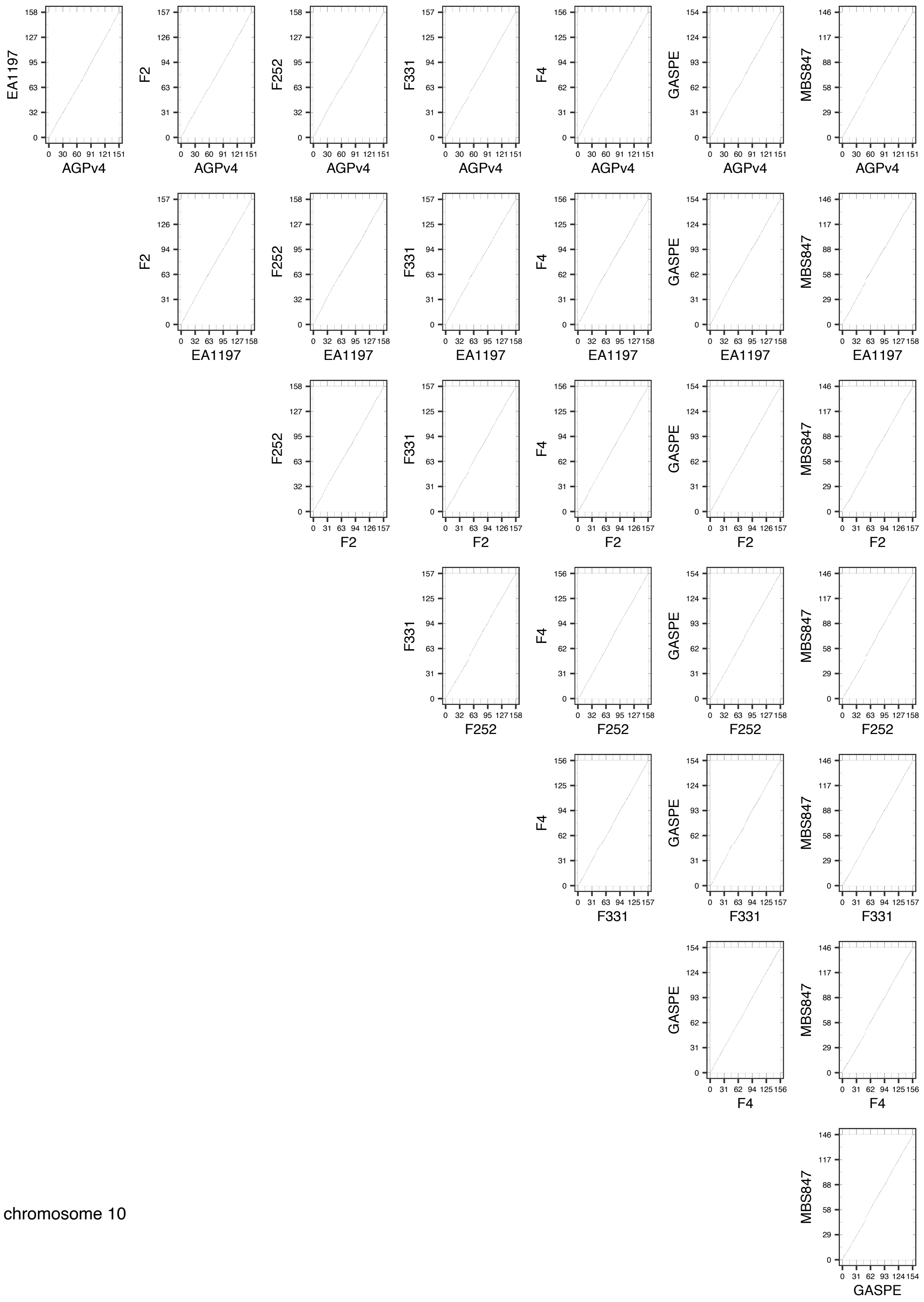


*Supplementary figure 1: Analysis of genome sequence contiguity by dot plot*

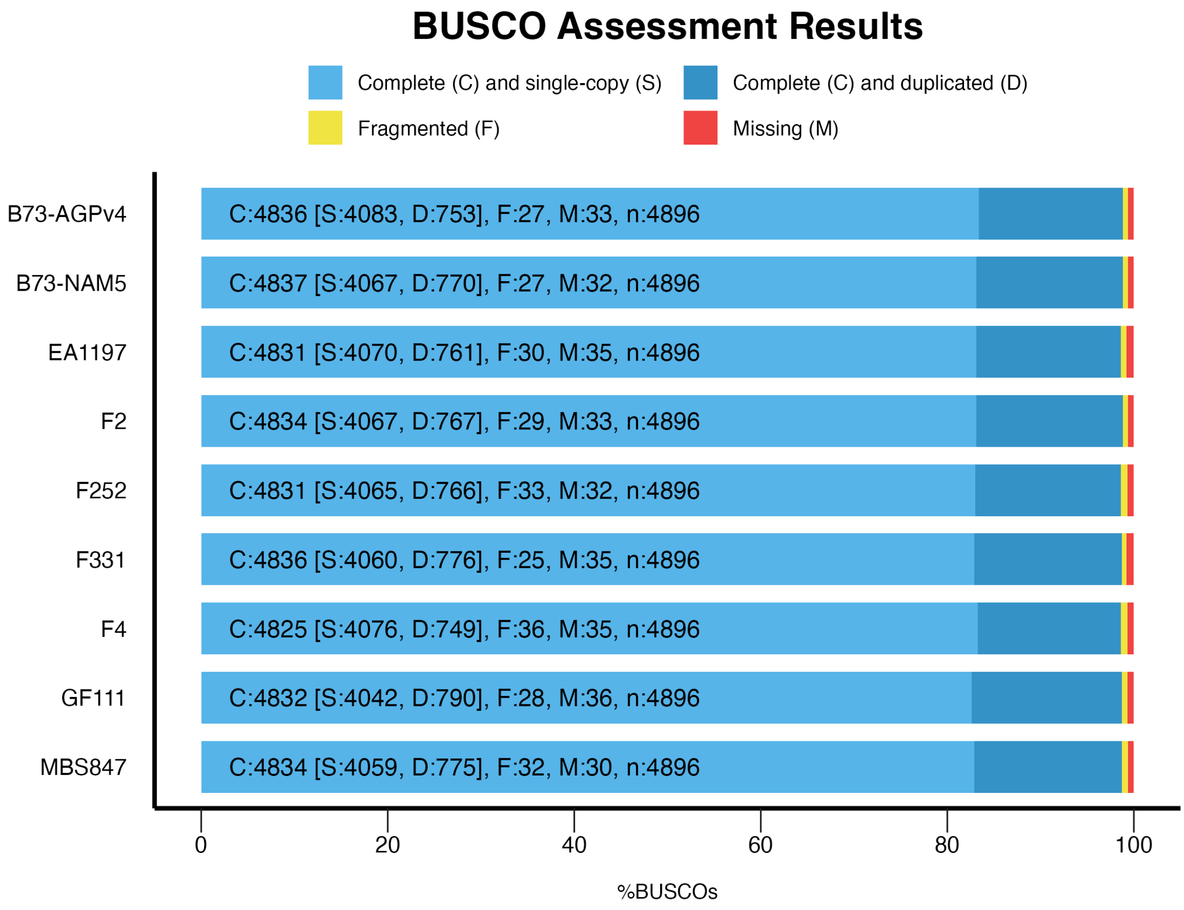


*Supplementary figure 2: BUSCO analysis of current assemblies and comparison with these of B73 AGPv4 and NAM5.*

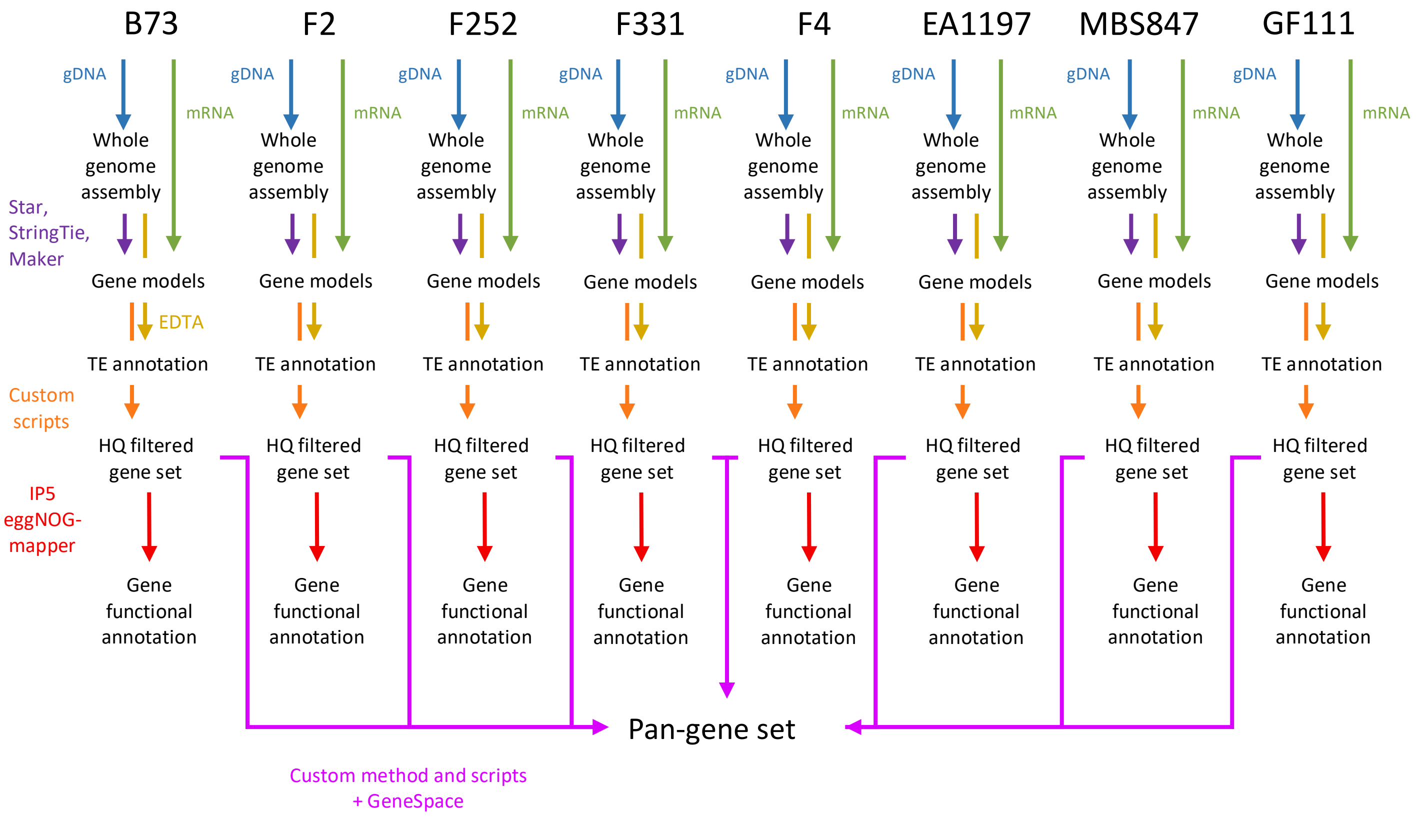


*Supplementary figure 3: Pan-gene set building workflow*

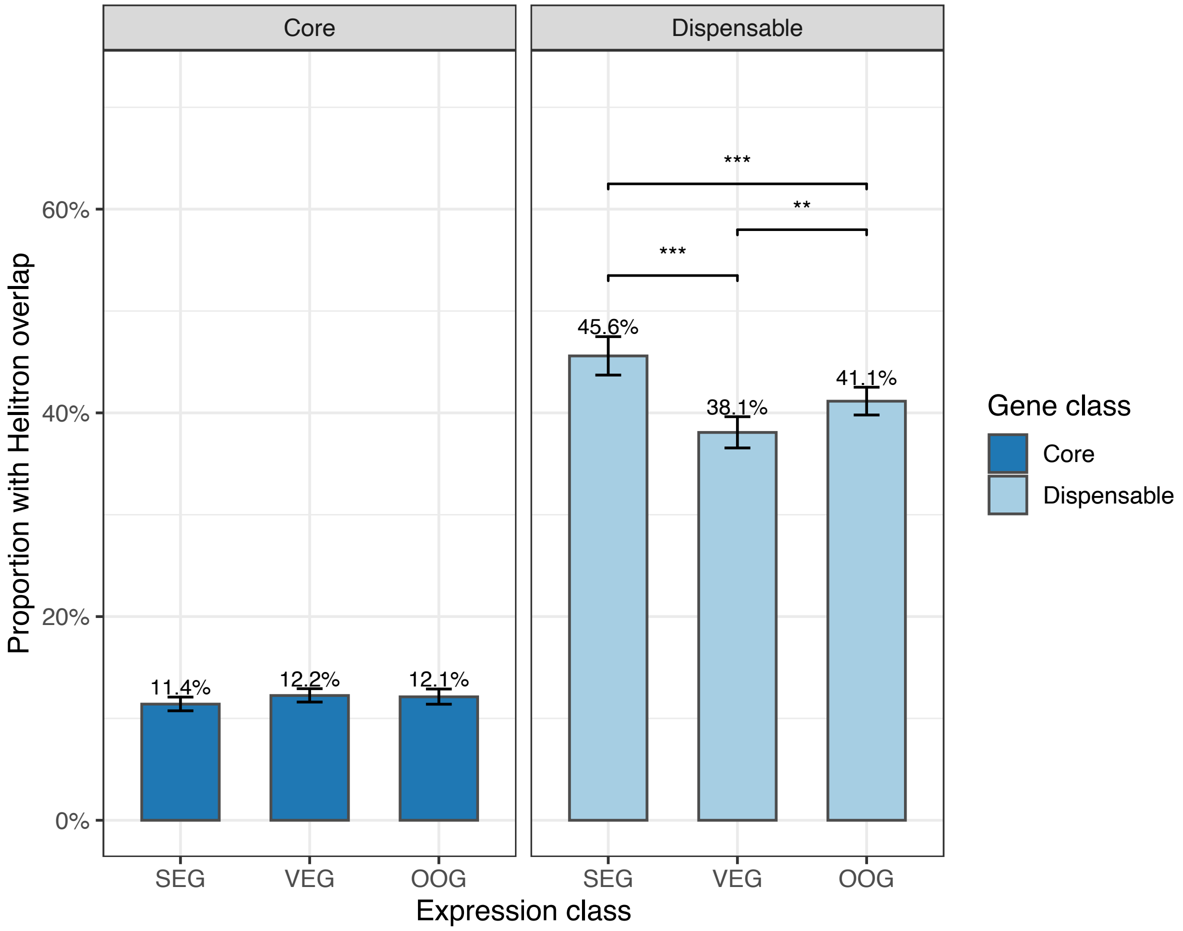


*Supplementary figure 4:* Helitron *transposable element overlap rate by expression class and gene class. Bars show the proportion of expressed genes (n = 37079) overlapping a* Helitron *TE within each combination of gene class (Core, n = 25549; Dispensable, n = 11530) and expression class (SEG: stably-expressed; VEG: variably-expressed; OOG: on-off genes). Bar colours indicate gene class (dark blue = Core, light blue = Dispensable). Error bars represent 95% Wilson confidence intervals. Significance brackets above bars indicate FDR-adjusted pairwise Fisher exact tests within each gene class: *** $p < 0.001$, ** $p < 0.01$, * $p < 0.05$; non-significant comparisons are omitted. The overall chi-square test across the six groups was significant (X2 = 4143.09, df = 5, $p < 2.2e-16$, Cramer's V = 0.334).*

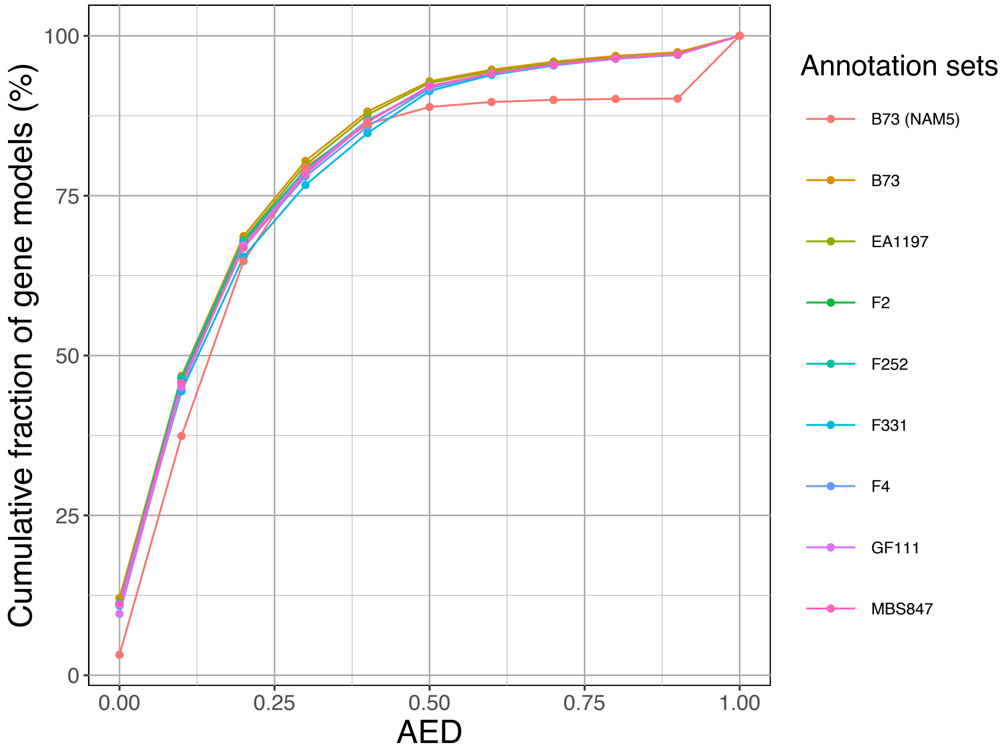


*Supplementary figure 5: AED score profile of each genotype predicted gene sets. B73 (NAM5) is used as a reference.*

### Supplementary tables

| Inbred line | Group | Sub-Group | Pedigree | Obtentor or Developer | Accession Code | Seedlot used |
|---|---|---|---|---|---|---|
| EA1197 | Tropical | Tropical Spanish | Mollar Almeria (Spain) | CSIC | EM1197_inra | EM1197_inra_SMH2009 |
| F2 | Flint | European Flint | Lacaune (France) | INRAE | FV2_inra | FV2_inra_SMH2014 |
| F252 | Dent | Lancaster / Iodent | CO125 | INRAE | FV252_inra | FV252_inra_MLN14 |
| F331 | Tropical | Tropical Highland | POB 86 (CIMMYT) | INRAE | FV331_inra | FV331_inra_SMH2004 |
| F4 | Flint | Northern Flint | Etoile de Normandie (France) | INRAE | FV4_inra | FV4_inra_SMH2005 |
| GF111 | Flint | Northern Flint | Gaspe (Canada) | University of Bologna | GF111_unibo | GF111_unibo_UBO2022 |
| MBS847 | Dent | Iodent | Mixed Dent | Mike Brayton Seeds | MBS847_inra | MBS847_inra_MLN14 |

*Supplementary table 1: Maize lines description*

| Genotype | Mapped reads (%) | Properly mapped reads (%) |
|---|---|---|
| B73 (AGPv4) | 97.16 | 95.27 |
| F2 | 99.67 | 89.5 |
| F252 | 99.39 | 88.79 |
| F331 | 99.70 | 93.35 |
| F4 | 99.58 | 92.27 |
| GF111 | 99.62 | 91.12 |
| EA1197 | 99.33 | 89.15 |
| MBS847 | 99.40 | 87.21 |

*Supplementary table 2: Fraction reads that align to the genome sequence of their genotype.*

| Genotype | Genes (total) | Core | Dispensable | Missing |
|---|---|---|---|---|
| B73 | 63606 | 46163 | 17443 | 12752 |
| EA1197 | 60783 | 46163 | 14620 | 15575 |
| F2 | 61344 | 46163 | 15181 | 15014 |
| F252 | 61334 | 46163 | 15171 | 15024 |
| F331 | 60810 | 46163 | 14647 | 15548 |
| F4 | 61360 | 46163 | 15197 | 14998 |
| GF111 | 61529 | 46163 | 15366 | 14829 |
| MBS847 | 61645 | 46163 | 15482 | 14713 |

*Supplementary table 3: Gene counts in each genotype*

| Terms | Df | Deviance | AIC | LRT | Pr(>Chi) |
|---|---|---|---|---|---|
| None | | 43951 | 56544 | | |
| Expression level | 1 | 42074 | 54670 | 1876.05 | < 2.2e-16 *** |
| Gene size | 1 | 43308 | 55903 | 643.00 | < 2.2e-16 *** |
| piN/piS | 1 | 43326 | 55922 | 624.13 | < 2.2e-16 *** |
| ENC | 1 | 43488 | 56083 | 462.65 | < 2.2e-16 *** |

*Supplementary table 4: Models comparison with single term addition showed highly significant Likelihood Ratio Test for each term. Null model: Core vs. dispensable ~ 1, Signif. codes: 0 '***' 0.001 '**' 0.01 '*' 0.05 '.' 0.1 ' ' 1.*

| Genotype | Tissue | Condition | Training set | Test set | ROC-AUC | AIC | Weight |
|---|---|---|---|---|---|---|---|
| B73 | DAP12_S | WW | 17866 | 4466 | 0.70 | 41963.95 | 24 |
| B73 | DAP35_S | WW | 17866 | 4466 | 0.68 | 41983.68 | 24 |
| B73 | DAS4_HY | WW | 17866 | 4466 | 0.75 | 41620.07 | 24 |
| B73 | DAS4_R | WW | 17866 | 4466 | 0.71 | 41698.55 | 24 |
| B73 | PS_EAR | WW | 17866 | 4466 | 0.69 | 41424.96 | 24 |
| B73 | PS_EAR | WD | 17866 | 4466 | 0.71 | 40178.96 | 22 |
| B73 | PS_IN1 | WW | 17866 | 4466 | 0.68 | 40548.30 | 22 |
| B73 | PS_IN1 | WD | 17866 | 4466 | 0.70 | 40515.42 | 22 |
| B73 | PS_LF | WW | 17866 | 4466 | 0.68 | 40776.64 | 22 |
| B73 | PS_LF | WD | 17866 | 4466 | 0.69 | 40380.02 | 22 |
| B73 | PS_SILK | WW | 17866 | 4466 | 0.72 | 40639.67 | 22 |
| B73 | PS_SILK | WD | 17866 | 4466 | 0.72 | 40469.33 | 22 |
| B73 | PS_TAS | WW | 17866 | 4466 | 0.68 | 42145.50 | 24 |
| B73 | PS_TAS | WD | 17866 | 4466 | 0.70 | 40585.68 | 22 |
| B73 | PTI_LF | WW | 17866 | 4466 | 0.67 | 42446.32 | 24 |
| B73 | PTI_LFI | WW | 17866 | 4466 | 0.73 | 41799.17 | 24 |
| B73 | V2_LF | WW | 17866 | 4466 | 0.71 | 40575.11 | 22 |
| B73 | V2_LFI | WW | 17866 | 4466 | 0.73 | 41534.56 | 24 |
| EA1197 | DAP12_S | WW | 17669 | 4417 | 0.71 | 41637.02 | 22 |
| EA1197 | DAP35_S | WW | 17669 | 4417 | 0.71 | 39908.11 | 20 |
| EA1197 | DAS4_HY | WW | 17669 | 4417 | 0.74 | 40972.69 | 22 |
| EA1197 | DAS4_R | WW | 17669 | 4417 | 0.70 | 41003.97 | 22 |
| EA1197 | PS_EAR | WW | 17669 | 4417 | 0.69 | 41248.78 | 22 |
| EA1197 | PS_EAR | WD | 17669 | 4417 | 0.70 | 41692.62 | 22 |
| EA1197 | PS_IN1 | WW | 17669 | 4417 | 0.70 | 40270.90 | 20 |
| EA1197 | PS_IN1 | WD | 17669 | 4417 | 0.69 | 40142.86 | 20 |
| EA1197 | PS_LF | WW | 17669 | 4417 | 0.65 | 39789.94 | 20 |
| EA1197 | PS_LF | WD | 17669 | 4417 | 0.69 | 40307.56 | 20 |
| EA1197 | PS_SILK | WW | 17669 | 4417 | 0.70 | 41529.94 | 22 |
| EA1197 | PS_SILK | WD | 17669 | 4417 | 0.68 | 41288.20 | 22 |
| EA1197 | PS_TAS | WW | 17669 | 4417 | 0.70 | 39695.11 | 20 |
| EA1197 | PS_TAS | WD | 17669 | 4417 | 0.69 | 39876.87 | 20 |
| EA1197 | PTI_LF | WW | 17669 | 4417 | 0.71 | 40473.80 | 20 |
| EA1197 | PTI_LFI | WW | 17669 | 4417 | 0.68 | 40769.50 | 22 |
| EA1197 | V2_LF | WW | 17669 | 4417 | 0.68 | 42033.46 | 22 |
| EA1197 | V2_LFI | WW | 17669 | 4417 | 0.72 | 41508.41 | 22 |
| F2 | DAP12_S | WW | 18112 | 4527 | 0.70 | 41529.33 | 22 |
| F2 | DAP35_S | WW | 18112 | 4527 | 0.72 | 40492.29 | 20 |
| F2 | DAS4_HY | WW | 18112 | 4527 | 0.72 | 41011.67 | 22 |
| F2 | DAS4_R | WW | 18112 | 4527 | 0.73 | 41992.21 | 22 |
| F2 | PS_EAR | WW | 18112 | 4527 | 0.76 | 40930.02 | 20 |
| F2 | PS_EAR | WD | 18112 | 4527 | 0.74 | 42211.45 | 22 |

| | | | | | | | |
|---|---|---|---|---|---|---|---|
| F2 | PS_IN1 | WW | 18112 | 4527 | 0.67 | 42296.54 | 22 |
| F2 | PS_IN1 | WD | 18112 | 4527 | 0.70 | 42396.75 | 22 |
| F2 | PS_LF | WW | 18112 | 4527 | 0.74 | 40690.98 | 20 |
| F2 | PS_LF | WD | 18112 | 4527 | 0.70 | 42562.17 | 22 |
| F2 | PS_SILK | WW | 18112 | 4527 | 0.72 | 40746.63 | 20 |
| F2 | PS_SILK | WD | 18112 | 4527 | 0.72 | 41829.26 | 22 |
| F2 | PS_TAS | WW | 18112 | 4527 | 0.71 | 40586.70 | 20 |
| F2 | PS_TAS | WD | 18112 | 4527 | 0.74 | 42424.03 | 22 |
| F2 | PTI_LF | WW | 18112 | 4527 | 0.73 | 39264.24 | 18 |
| F2 | PTI_LFI | WW | 18112 | 4527 | 0.70 | 42021.38 | 22 |
| F2 | V2_LF | WW | 18112 | 4527 | 0.71 | 42678.57 | 22 |
| F2 | V2_LFI | WW | 18112 | 4527 | 0.72 | 42124.22 | 22 |
| F252 | DAP12_S | WW | 17564 | 4390 | 0.69 | 40617.98 | 22 |
| F252 | DAP35_S | WW | 17564 | 4390 | 0.70 | 40813.60 | 22 |
| F252 | DAS4_HY | WW | 17564 | 4390 | 0.74 | 40003.06 | 22 |
| F252 | DAS4_R | WW | 17564 | 4390 | 0.69 | 39946.83 | 22 |
| F252 | PS_EAR | WW | 17564 | 4390 | 0.68 | 40599.51 | 22 |
| F252 | PS_EAR | WD | 17564 | 4390 | 0.71 | 40521.97 | 22 |
| F252 | PS_IN1 | WW | 17564 | 4390 | 0.67 | 40924.85 | 22 |
| F252 | PS_IN1 | WD | 17564 | 4390 | 0.67 | 40853.56 | 22 |
| F252 | PS_LF | WW | 17564 | 4390 | 0.68 | 40899.90 | 22 |
| F252 | PS_LF | WD | 17564 | 4390 | 0.68 | 38550.36 | 20 |
| F252 | PS_SILK | WW | 17564 | 4390 | 0.69 | 38989.21 | 20 |
| F252 | PS_SILK | WD | 17564 | 4390 | 0.67 | 40569.94 | 22 |
| F252 | PS_TAS | WW | 17564 | 4390 | 0.71 | 39118.27 | 20 |
| F252 | PS_TAS | WD | 17564 | 4390 | 0.69 | 40472.67 | 22 |
| F252 | PTI_LF | WW | 17564 | 4390 | 0.67 | 39046.95 | 20 |
| F252 | PTI_LFI | WW | 17564 | 4390 | 0.67 | 40116.68 | 22 |
| F252 | V2_LF | WW | 17564 | 4390 | 0.68 | 39202.22 | 20 |
| F252 | V2_LFI | WW | 17564 | 4390 | 0.72 | 40196.03 | 22 |
| F331 | DAP12_S | WW | 17504 | 4375 | 0.70 | 39512.22 | 22 |
| F331 | DAP35_S | WW | 17504 | 4375 | 0.68 | 43104.12 | 26 |
| F331 | DAS4_HY | WW | 17504 | 4375 | 0.72 | 39085.49 | 22 |
| F331 | DAS4_R | WW | 17504 | 4375 | 0.67 | 40920.84 | 24 |
| F331 | PS_EAR | WW | 17504 | 4375 | 0.68 | 39252.02 | 22 |
| F331 | PS_EAR | WD | 17504 | 4375 | 0.67 | 41645.31 | 24 |
| F331 | PS_IN1 | WW | 17504 | 4375 | 0.68 | 42097.05 | 24 |
| F331 | PS_IN1 | WD | 17504 | 4375 | 0.70 | 39717.13 | 22 |
| F331 | PS_LF | WW | 17504 | 4375 | 0.71 | 38373.78 | 20 |
| F331 | PS_LF | WD | 17504 | 4375 | 0.67 | 43269.56 | 26 |
| F331 | PS_SILK | WW | 17504 | 4375 | 0.68 | 40956.57 | 24 |
| F331 | PS_SILK | WD | 17504 | 4375 | 0.67 | 39634.58 | 22 |
| F331 | PS_TAS | WW | 17504 | 4375 | 0.66 | 40589.70 | 24 |
| F331 | PS_TAS | WD | 17504 | 4375 | 0.70 | 39372.31 | 22 |

| | | | | | | | |
|---|---|---|---|---|---|---|---|
| F331 | PTI_LF | WW | 17504 | 4375 | 0.65 | 41409.26 | 24 |
| F331 | PTI_LFI | WW | 17504 | 4375 | 0.68 | 40729.97 | 24 |
| F331 | V2_LF | WW | 17504 | 4375 | 0.70 | 39979.61 | 22 |
| F331 | V2_LFI | WW | 17504 | 4375 | 0.69 | 40565.06 | 24 |
| F4 | DAS4_HY | WW | 17043 | 4259 | 0.70 | 39742.36 | 24 |
| F4 | DAS4_R | WW | 17043 | 4259 | 0.68 | 39900.44 | 24 |
| F4 | PS_EAR | WW | 17043 | 4259 | 0.71 | 38663.50 | 22 |
| F4 | PS_EAR | WD | 17043 | 4259 | 0.68 | 38805.96 | 22 |
| F4 | PS_IN1 | WW | 17043 | 4259 | 0.65 | 40225.72 | 24 |
| F4 | PS_IN1 | WD | 17043 | 4259 | 0.68 | 38669.54 | 22 |
| F4 | PS_LF | WW | 17043 | 4259 | 0.67 | 39111.26 | 22 |
| F4 | PS_LF | WD | 17043 | 4259 | 0.69 | 39176.76 | 22 |
| F4 | PS_SILK | WW | 17043 | 4259 | 0.69 | 36987.95 | 20 |
| F4 | PS_SILK | WD | 17043 | 4259 | 0.68 | 38883.19 | 22 |
| F4 | PS_TAS | WW | 17043 | 4259 | 0.70 | 38884.37 | 22 |
| F4 | PS_TAS | WD | 17043 | 4259 | 0.68 | 38831.61 | 22 |
| F4 | PTI_LF | WW | 17043 | 4259 | 0.68 | 38890.36 | 22 |
| F4 | PTI_LFI | WW | 17043 | 4259 | 0.65 | 39472.54 | 24 |
| F4 | V2_LF | WW | 17043 | 4259 | 0.66 | 40335.53 | 24 |
| F4 | V2_LFI | WW | 17043 | 4259 | 0.70 | 40136.44 | 24 |
| GF111 | DAS4_R | WW | 17515 | 4378 | 0.72 | 40276.07 | 22 |
| GF111 | PTI_LFI | WW | 17515 | 4378 | 0.73 | 38521.60 | 20 |
| GF111 | PS_TAS | WW | 17515 | 4378 | 0.70 | 40429.66 | 22 |
| GF111 | V2_LFI | WW | 17515 | 4378 | 0.70 | 40123.79 | 22 |
| GF111 | PS_IN1 | WW | 17515 | 4378 | 0.69 | 38910.18 | 20 |
| GF111 | V2_LF | WW | 17515 | 4378 | 0.70 | 39465.05 | 20 |
| GF111 | PS_EAR | WD | 17515 | 4378 | 0.68 | 40699.65 | 22 |
| GF111 | PTI_LF | WW | 17515 | 4378 | 0.69 | 39253.17 | 20 |
| GF111 | DAP12_S | WW | 17515 | 4378 | 0.70 | 38834.18 | 20 |
| GF111 | PS_IN1 | WD | 17515 | 4378 | 0.69 | 39225.08 | 20 |
| GF111 | DAP35_S | WW | 17515 | 4378 | 0.71 | 39245.75 | 20 |
| GF111 | PS_EAR | WW | 17515 | 4378 | 0.67 | 40305.82 | 22 |
| GF111 | PS_SILK | WD | 17515 | 4378 | 0.67 | 40872.64 | 22 |
| GF111 | PS_LF | WW | 17515 | 4378 | 0.66 | 39194.87 | 20 |
| GF111 | PS_TAS | WD | 17515 | 4378 | 0.69 | 40849.53 | 22 |
| GF111 | DAS4_HY | WW | 17515 | 4378 | 0.68 | 41715.08 | 24 |
| GF111 | PS_SILK | WW | 17515 | 4378 | 0.67 | 40909.41 | 22 |
| MBS847 | DAP12_S | WW | 17347 | 4336 | 0.69 | 38547.66 | 20 |
| MBS847 | DAP35_S | WW | 17347 | 4336 | 0.69 | 40661.40 | 22 |
| MBS847 | DAS4_HY | WW | 17347 | 4336 | 0.72 | 40134.05 | 22 |
| MBS847 | DAS4_R | WW | 17347 | 4336 | 0.70 | 40447.37 | 22 |
| MBS847 | PS_EAR | WW | 17347 | 4336 | 0.66 | 40227.92 | 22 |
| MBS847 | PS_EAR | WD | 17347 | 4336 | 0.69 | 38507.12 | 20 |
| MBS847 | PS_IN1 | WW | 17347 | 4336 | 0.65 | 40620.11 | 22 |

| | | | | | | | |
|---|---|---|---|---|---|---|---|
| MBS847 | PS_IN1 | WD | 17347 | 4336 | 0.67 | 40663.27 | 22 |
| MBS847 | PS_LF | WW | 17347 | 4336 | 0.68 | 41289.89 | 22 |
| MBS847 | PS_LF | WD | 17347 | 4336 | 0.66 | 38898.18 | 20 |
| MBS847 | PS_SILK | WW | 17347 | 4336 | 0.69 | 38895.72 | 20 |
| MBS847 | PS_TAS | WW | 17347 | 4336 | 0.65 | 40382.33 | 22 |
| MBS847 | PS_TAS | WD | 17347 | 4336 | 0.74 | 39435.11 | 20 |
| MBS847 | PTI_LF | WW | 17347 | 4336 | 0.67 | 39578.61 | 20 |
| MBS847 | PTI_LFI | WW | 17347 | 4336 | 0.68 | 40430.78 | 22 |
| MBS847 | V2_LF | WW | 17347 | 4336 | 0.68 | 38954.89 | 20 |
| MBS847 | V2_LFI | WW | 17347 | 4336 | 0.70 | 40171.51 | 22 |
| | | | | Average | 0.69 | | |
| | | | | SD | 0.02 | | |

*Supplementary table 5: GLM for prediction of core versus dispensable status of gene from various gene properties (core* vs. *dispensable ~ average expression level + CDS size + piN/piS + GC3%). A glm was trained for each experiment of gene expression level measurements: genotype, tissue and condition. GLM were weighted to address unbalanced proportion of core and dispensable genes in training and test sets. Size of training and sample sets are reported.*

| Organ group | Tissue description | Growth stage at collection | Growth method | Tissue designation | ***short name*** | Conditions | No. of plants per replicate |
|---|---|---|---|---|---|---|---|
| **Root** | Pooled roots (from emerging point to root tip) | 4 DAS | Perlite, without direct light | DAS4_R | *r* | WW | 3 |
| **Leaf** | Hypocotyl (from emerging point to 5 mm above meristematic ring) | 4 DAS | Perlite, without direct light | DAS4_HY | *h* | WW | 3 |
| | Pooled immature leaves (white parts of leaf 3 and onwards) | V2 | Greenhouse | V2_LFI | *pc* | WW | 1 |
| | Pooled leaf blades (green expanded parts of leaves 1 to 4) | V2 | Greenhouse | V2_LF | *pm* | WW | 1 |
| | Last visible leaf: immature part (white part of leaf covered by other leaves) | Post-Tassel Initiation | Greenhouse | PTI_LFI | *fc* | WW | 1 |
| | Last ligulated leaf: middle part of leaf blade, without midrib | Post-Tassel Initiation | Greenhouse | PTI_LF | *fm* | WW | 1 |
| | Last ligulated leaf below flag leaf: middle part of leaf blade, without midrib | Pollen shed | Greenhouse | PS_LF | *f* | WW and WD | 1 |
| **Internode** | Second internode below ear (1/3rd of internode immediately above node) | Pollen shed | Greenhouse | PS_IN0 | *e0* | WW and WD | 1 |
| | First internode below ear (1/3rd of internode immediately above node) | Pollen shed | Greenhouse | PS_IN1 | *e1* | WW and WD | 1 |
| | Ear internode (1/3rd of internode immediately above node) | Pollen shed | Greenhouse | PS_IN2 | *e2* | WW and WD | 1 |
| **Reproductive** | Pooled silks, growing parts under husk | Pollen shed | Greenhouse | PS_SILK | *s* | WW and WD | 1 |
| | Tassel, pool of parts harbouring fresh pollen | Pollen shed | Greenhouse | PS_TAS | *p* | WW and WD | 1 |
| | Pre-pollination ear, middle part (1/3rd top part and 1/5th bottom part removed) | Pollen shed | Greenhouse | PS_EAR | *e* | WW and WD | 1 |
| **Seed** | Whole seed (embryo, endosperm and pericarp), end of early development stage | 12 DAP | Field (Summer 2017, South West France) | DAP12_S | *g12* | WW | 1 |
| | Whole seed (embryo, endosperm and pericarp), late filling stage | 35 DAP | Field (Summer 2017, South West France) | DAP35_S | *g15* | WW | 1 |

DAS: days after sowing; V*n:* vegetative stage corresponding to the number *n* of ligulated (fully expanded) leaves; PS: pollen shed; DAP: days after pollination

*Supplementary table 6: Description of samples used for RNA-seq analyses.*
